# *Trypanosoma brucei* ultra-structure cell modifications caused by knockdown of the ribonuclease Rrp44/Dis3

**DOI:** 10.1101/2020.02.29.971424

**Authors:** Giovanna Cesaro, Priscila M. Hiraiwa, Flavia R. G. Carneiro, Valérie Rouam, Pierre Legrand, José Javier Conesa, Matthieu Réfrégiers, Eva Pereiro, Beatriz G. Guimarães, Frédéric Jamme, Nilson I. T. Zanchin

## Abstract

Rrp44/Dis3 is an essential protein conserved in all eukaryotes that functions in the maturation of many different RNA precursors and RNA surveillance. The *Trypanosoma brucei* Rrp44/Dis3 homologue (TbRRP44) is required for maturation of pre-rRNA, spliced leader, and U3 snoRNA precursors. Its depletion leads to inhibition of cell proliferation and eventually to cell death. In this work, we showed that TbRRP44 depletion causes a massive expansion of acidic and lysosome-derived vacuoles, enlargement of cell and nuclei sizes without changes in DNA content, mitochondrial inactivation, and autophagy induction. Consistently, 3D reconstructions using cryo-soft X-ray tomography revealed extreme vacuolation of the cytoplasm and numerous cellular alterations, including an increase in size and number of calcium-containing vesicles and lipid droplets. These multiple defects indicate that a combination of alterations converge to induce lysosome expansion. With time, the cytoplasm is taken up by lysosome-derived vacuoles, which may be a final stage leading to the cell death triggered by TbRRP44 depletion. These studies provide the first evidence on the ultra-structure cell modifications caused by deficiency of this essential ribonuclease in *T. brucei*.

## Introduction

The first Rrp44/Dis3 homologue was isolated from the fission yeast *Schizosaccharomyces pombe* by complementation of the “defective in sister chromatid disjoining” 3 *(dis3*) mutant, a conditional cold-sensitive strain which did not accomplish mitotic chromosome separation during cell division when incubated at the non-permissive temperature ^1^. The *S. pombe dis3*^+^ gene was subsequently shown to encode an essential 110 kDa protein required for mitotic cell division ^2^. The biochemical function of the protein encoded by *dis3*^+^ homologues was revealed later when the RNA exosome complex was described ^3^. Due to its role in RNA processing, the *Saccharomyces cerevisiae* homologue was named RNA processing protein 44 (Rrp44). Following the guidelines for the genetic nomenclature for *Trypanosoma* and *Leishmania* ^4^, the *T. brucei* Rrp44/Dis3 homologue will be referred to as TbRRP44 in this paper.

*S. cerevisiae* and human Rrp44/Dis3 present 3’-5’ exonucleolytic activity as well as endonucleolytic activity carried out by RNB and PIN domains, respectively ^5–8^. The activity of the RNB domain is essential for cell viability while the PIN domain contributes to Rrp44/Dis3 function but seems not to be essential ^5–10^. Rrp44/Dis3 participates in the maturation and degradation of multiple RNA types ^11–16^ and functions in association with the RNA exosome complex, being one of its catalytical subunits together with Rrp6, except for the trypanosomatid homologue, which was not found in association with the exosome complex ^17,18^. Nevertheless, the knockdown of structural exosome subunits produced similar phenotypes on the maturation of the *T. brucei* 5.8S rRNA 3’-end as the knockdown of TbRRP44 ^19^. This indicates that even if not in physical association, it must cooperate with the exosome at least for maturation of the 5.8S rRNA 3’-end. In a previous study, we have shown that like the *S. cerevisiae* and human homologues, the exonuclease activity of TbRRP44 is essential for *T. brucei* viability ^20^. While the physical presence of the PIN domain is required for *T. brucei* viability, its catalytic activity is not ^20^. TbRRP44 can degrade non-structured and structured RNA substrates in vitro ^21^ and is required for maturation of small and large subunit pre-rRNAs, spliced leader and snoRNA precursors, indicating that it plays a general role in RNA processing also in *T. brucei* ^20^.

Rrp44/Dis3 homologues were shown to play essential functions in early developmental stages of multicellular model organisms. Knockdown of Rrp44//Dis3 inhibits female gametophyte development and early embryogenesis in *Arabidopsis thaliana* ^22,23^ and, inhibits growth and cause death at the larval stage in *Drosophila melanogaster* ^24^ and in *Caenorhabditis elegans* ^25^. Homozygous Rrp44/Dis3 knockout mice embryos show overaccumulation of the homeobox transcription factor *Pou6f1* mRNA and arrest at the morula stage ^26^. Although intriguing but consistent with the initial findings on the requirement of Rrp44/Dis3 for mitotic chromosome segregation, several studies have supported a functional interplay between Rrp44/Dis3 function and genome stability. *S. pombe* Dis3 is required for kinetochore formation and kinetochore-microtubule interactions ^27^. Mutations in the exonuclease domain of *S. cerevisiae* Rrp44p lead to genome instability ^28^. The phenotypes observed for Rrp44/Dis3 deficiency in *D. melanogaster* also included delayed mitosis, aneuploidy, and overcondensed chromosomes ^25^. In the case of mammalian cells, Rrp44/Dis3 has been implicated in the control of chromosomal architecture during B cell development ^29^, being required for V(D)J recombination at the immunoglobulin heavy chain (*Igh*) locus ^29,30^. Activated B cells deficient in Rrp44/Dis3 show increased DNA:RNA hybrids in the V(D)J regions ^29^. Moreover, the regulation of B cell development by the RNA exosome has been proposed to take place by a noncoding RNA processing mechanism ^30^. Rrp44/Dis3, along with other exosome subunits, were shown to play an essential role in the balance between proliferation and differentiation of erythroid cells. Rrp44/Dis3 deficiency in mice primary erythroid precursors induces erythroid cell maturation and blocks proliferation ^31^ leading to an exhaustion of erythroid progenitor cells ^32^.

The phenotypes related to genome function, stability, and organization under conditions of Rrp44/Dis3 deficiency may result from its role in RNA surveillance, which comprises a series of mechanisms responsible for maintaining the balance of both coding and non-coding RNA molecules along the development ^13,33–36^. This is supported by the fact that Rrp44/Dis3 deficiency causes a general accumulation of DNA:RNA hybrids, which results in increased mutational rate and genome instability ^37^. Furthermore, depletion of Rrp44/Dis3 in human colon carcinoma derived HCT116 cells leads to accumulation of enhancer RNAs, promoter upstream transcripts (PROMPTs), premature cleavage, and polyadenylation products ^38^.

Alterations in RNA surveillance pathways due to Rrp44/Dis3 deficiency may result in diverse biological outcomes, depending on the cell type and the nature of Rrp44/Dis3 mutations. In some cases, Rrp44/Dis3 deficiency can result in enhancement of cell proliferation. For instance, global genome and gene expression analyses have established a connection between defective mutations in Rrp44/Dis3 with multiple myeloma development ^39,40^. RNAi knockdown of Rrp44/Dis3 in tissue culture impairs maturation of let-7 miRNAs leading to an increase of the pluripotency factor LIN28B, which results in upregulation of growth-promoting oncogenic proteins such as MYC and RAS ^41^. Rrp44/Dis3 deficiency has been linked to the promotion of cell proliferation also in *C. elegans*, *D. melanogaster* eggs, and mouse B cells stimulated with bacterial lipopolysaccharides and interleukin 4 ^25^.

During previous studies to characterize the role of TbRRP44 in RNA processing^20,21,42^, we have observed that the knockdown of TbRRP44 causes cellular defects in *T. brucei* procyclic cells, which have not been described previously. Therefore, in the present work, we have investigated the cellular alterations that take place as a consequence of the absence of the TbRRP44 in the cell. This work complements the previous data by providing a description of the defects that lead to *T. brucei* cell death upon depletion of this essential protein. Since the genetic and immunological tools equivalent to the ones used for the characterization of nucleolar stress in *S. cerevisiae* and mammalian cells are not currently available for *T. brucei*, we combined different analyses using cryo-soft X-ray tomography (cryo-SXT) and fluorescence imaging methods to investigate these phenotype switches. Biochemical and molecular markers were used to obtain information on the organelles affected. We show that depletion of TbRRP44 blocked proliferation in parallel with general alterations of the cells. 3D reconstructions revealed an extreme increase in the size of low absorbance vacuoles and a general enlargement of various types of cellular vesicles. Compared to other studies that have described *T. brucei* cell death ^43–46^, the cellular defects caused by TbRRP44 depletion are unique in the sense that they converge to cause an expansion of lysosome derived vesicles that ultimately take up most of the cytoplasm. This work reveals unprecedented details of the cellular changes that occur in *T. brucei* cells on the route to death after the depletion of this essential protein.

## Results

### TbRRP44 knockdown results in activation autophagy response

Knockdown of TbRRP44 impairs large ribosomal subunit RNA maturation, causing accumulation of unprocessed pre-rRNA, and disrupting ribosome synthesis ^19,20,42^. In animal cells, similar defects caused by mutation or deficiency in genes encoding ribosomal proteins and ribosome synthesis factors result in nucleolar stress leading to activation of autophagy, cycle arrest and/or apoptosis ^47,48^. Autophagy starts with the formation of autophagosomes, which are dense double membrane structures that later fuse with lysosomes forming autophagolysosomes, and can be monitored by using autophagy markers ^49,50^. To determine whether there was a connection between TbRRP44 deficiency and autophagy activation, the TbRRP44 conditional strain was transformed with a plasmid expressing the autophagy marker ATG8.2 fused to the yellow fluorescent protein ^51^.Autophagy induction was evaluated as described by Li and co-workers ^49^. The ratio of autophagosomes per cell was compared between control and TbRRP44 knockdown cells incubated either in SDM 79 medium or in PBS containing 1 g/L glucose (gPBS) for autophagy induction (Fig. 1A and B). While control cells require incubation in gPBS to develop autophagosomes, the TbRRP44 knockdown cells incubated in SDM 79 medium already present autophagosomes (Fig. 1B). The number of autophagosomes increases when the TbRRP44 knockdown cells are incubated in the gPBS solution. This result shows that TbRRP44 depletion results in induction of the autophagy response and indicates that autophagy participates in the events that lead to inhibition of proliferation of TbRRP44-depleted cells.

**Figure 1.**
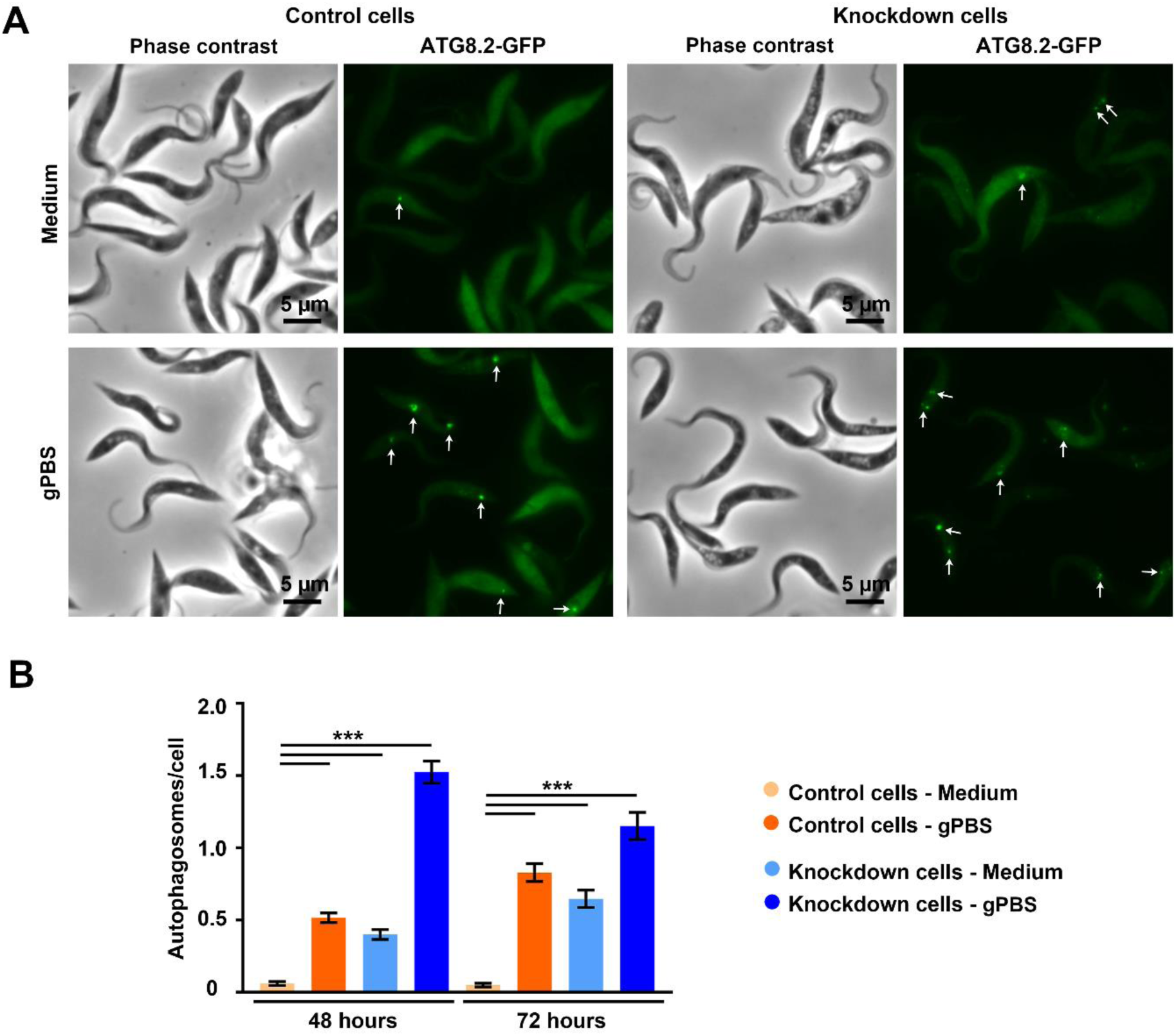
Autophagy response in TbRRP44 knockdown cells. (**A**) Representative images used in autophagosome quantification. Control and TbRRP44 knockdown cells were incubated under autophagosome induction conditions (gPBS: PBS containing 1 g/L glucose) or in SDM 79 culture medium. Left: phase contrast images. Right: autophagy marker ATG8.2 fused to YFP and immunodetected using an anti-GFP nanobody. White arrows indicate the autophagosomes. Scale bar, 5 µm. (**B**) Quantification of the number of autophagosomes per cell. The mean number of autophagosomes per cell of was determined by counting between 300 and 700 cells depending on the experimental condition. The exact number of cells counted for each experimental condition is given in the Materials and Methods section. Differences between samples was assessed by one-way analysis of variance (ANOVA) followed by Tukey’s multiple comparison tests using GraphPad Prism version 7.01 software. *: 0.01<p<0.05; **: 0.001<p<0.01; ***: p<0.001.

### Moderate changes in DNA content and cellular membrane integrity in TbRRP44 knockdown cells

In our assays, the number of cells in the TbRRP44 knockdown cultures remained nearly unchanged starting at 48 hours after RNAi induction, indicating that there is a stabilization between the rates of cell division and death. To evaluate if TbRRP44 deficiency leads to cell cycle defects, the DNA content of control and TbRRP44 knockdown cells was evaluated by cytometry. However, only moderate changes were observed at 48 and 72 hours of knockdown (Fig. 2A and B). At these times, there is a small accumulation of cells at the G1 stage with a small reduction of cells at the S and G2 stages. A stronger alteration in DNA content can be detected only after 96 hours of knockdown, when the sub-G1 population reached ∼30% with a parallel reduction of all other stages (Fig. 2A and B). The sub-G1 population comprises cells that are undergoing DNA damage ^52^, indicating that the DNA damage starts to appear after the cells stop dividing.

**Figure 2.**
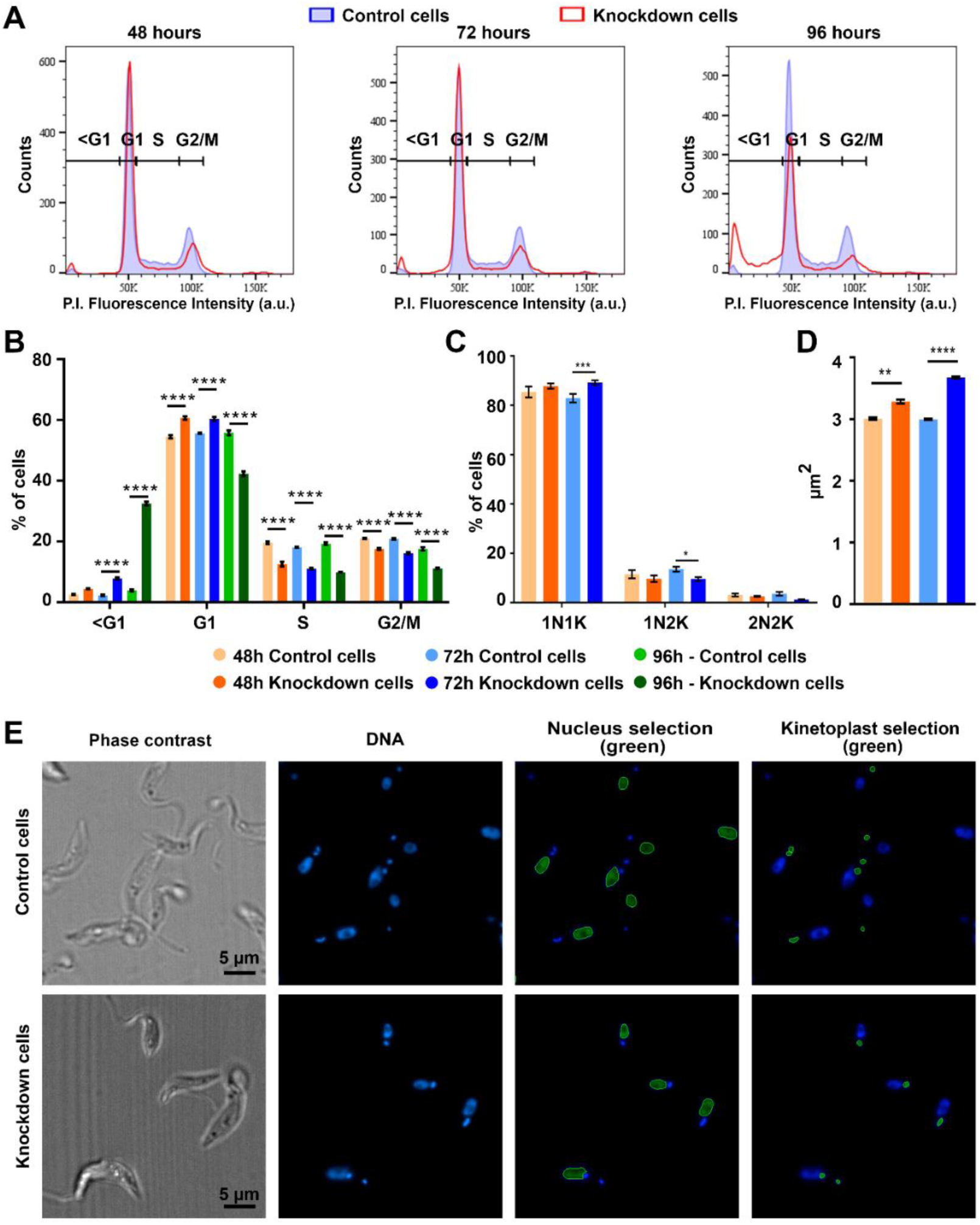
Changes in DNA content, nucleus/kinetoplast ratio and increase in nuclear area. For cell cycle analysis based on DNA content, cells were collected at 48, 72 and 96 hours after induction of RNAi for TbRRP44 knockdown. For quantification of nucleus/kinetoplast (N/K) ratio, cells were collected at 48 and 72 hours after RNAi induction. (**A**) Cell cycle profiles determined by flow cytometry cell sorting based on DNA staining with propidium iodide. (**B**) Percentage of cells in each phase of the cell cycle. At the earlier times, there is a small increase of TbRRP44-depleted cells in the G1 phase in parallel with a decrease of cells in the S and G2/M phases. At 96 hours, there is an increase in the sub-G1 population with a decrease of all other stages in TbRRP44-depleted cells, indicating that the cells start to undergo DNA damage. (**C**) Quantification of cells according to nucleus/kinetoplast ratio configuration. There is a small but significant increase of the 1N/1K population, accompanied by a decrease in the 1N/2K population in TbRRP44 knockdown cells at 72 hours of depletion. (**D**) Quantification of nuclei area showing higher average area for TbRRP44-depleted cells (**E**) Representative images used in the analysis of N/K ratio per cell. Nuclei and kinetoplasts were stained with DAPI and selected using Harmony 4.6 software from PerkinElmer. Differences between samples was assessed by one-way analysis of variance (ANOVA) followed by Tukey’s multiple comparison tests using GraphPad Prism version 7.01 software. *: 0.01<p<0.05; **: 0.001<p<0.01; ***: p<0.001; ****: p<0.0001.

In trypanosomatids, DNA replication and division of the kinetoplast (K) precedes nuclear (N) DNA replication and division so that the nuclear/kinetoplast ratio also can provide information on the cell cycle stages. Cells presenting 1N/1K comprise the G1 and S phases, cells presenting 1N/2K are in phase G2, and cells presenting 2N/2K are in the mitotic or post-mitotic phases ^53,54^. The N/K ratio of over one thousand cells per treatment was analyzed using a high content imaging system. Consistently with the cytometry data, only a moderate increase of 1N/1K cells was observed at 72 hours after TbRRP44 knockdown (Fig. 2C and E). During these analyses, a change was noticed in the nuclear size. By determining the nuclear area, we observed an increase of ∼10% and ∼20% at 48 and 72 hours after TbRRP44 depletion, respectively (Fig. 2D). Nuclear enlargement without an increase in DNA content has been related to senescence in mammalian cells in response to treatments with excess thymidine ^55^, resveratrol, quercetin ^56^, and sirtuin inhibitors ^57^. There is no similar phenotype reported for trypanosomatids, but nuclear enlargement may represent a senescence-like process that results from TbRRP44 deficiency.

In parallel with the cell cycle analysis, we have evaluated cell membrane asymmetry and integrity, which could reveal if the cells are entering a cell death process after stopping proliferating. Cellular pro-death signals usually trigger the inversion of phosphatidylserine residues, leading to loss of cell membrane asymmetry. In addition, cell membrane damages appear when cells start to undergo apoptosis or necrosis processes. The ratio of phosphatidylserine inversion was determined using annexin V conjugated with Alexa fluor 488 in the same cells stained in parallel with propidium iodide. Surprisingly, even at 96 hours of TbRRP44 knockdown, the ratio of cells positive for propidium iodide falls below 5%, showing that the cell membrane is not undergoing high damage and indicating that the cells are not undergoing a strong necrosis process (Fig. 3). The population of positive cells for annexin V ranged from ∼12% to 16% for the three time points of knockdown analyzed (Fig. 3). This indicates that loss of cell membrane asymmetry is taking place at a low rate in TbRRP44 knockdown cells. Similarly, only a fraction of approximately 10-12% of the cell population showed positive staining for both annexin V and propidium iodide in the 72- and 96-hour time points (Fig. 3). The low ratio of cell population positive for both markers indicates that the cell membrane remains relatively stable for long TbRRP44 depletion times.

**Figure 3.**
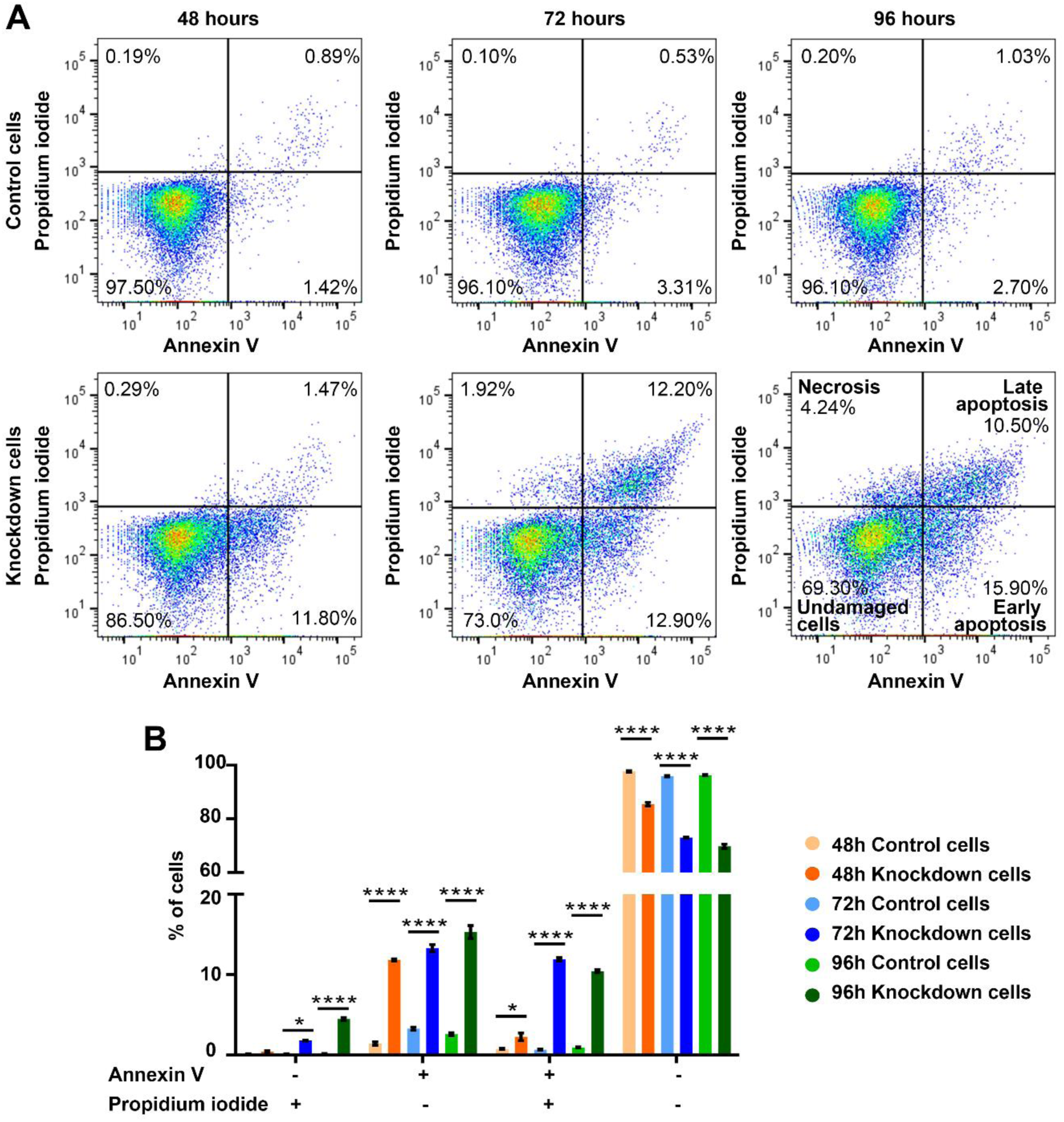
Quantification of pro-death and death events using membrane asymmetry and membrane damage markers. (**A**) Flow cytometry profiles of cells stained with annexin V-Alexa fluor 488 and propidium iodide. TbRRP44 knockdown cells were collected for analysis at 48, 72, and 96 hours after RNAi induction. Upper panel: control cells. Lower panel: TbRRP44 knockdown cells. (**B**) Graph showing the percentage of cells per category. Undamaged cells: cells negative for staining with both markers. Necrosis: cells positive for propidium iodide. Early apoptosis: cells positive for annexin V. Late apoptosis: cells positive for both markers. The combination of cells at early and late apoptosis stages reaches ∼30% of the cell population starting after 72 hours of TbRRP44 depletion, indicating relatively mild damages to the cell membrane.

### Cryo-soft X-ray tomography reveals extensive cellular ultrastructure alterations in TbRRP44 knockdown cells

The relatively mild damages observed in the cell membrane and nuclear DNA described above contrasted with our preliminary fluorescence microscopic analyses indicating that important alterations are taking place in the cytoplasm of TbRRP44 knockdown cells. This raised an intriguing question about what organelles are affected after the cells stop proliferating. In order to investigate this point, we have evaluated the ultrastructural alterations of these cells using cryo-soft X-ray tomography (cryo-SXT), which allows reconstitution of the 3D structure of whole cells at nanometer resolution ^58,59^. Cryo-SXT is usually performed with vitrified samples near native state. However, due to flagellar activity, the living *T. brucei* procyclic form presents high motility and does not adhere well to poly-L-lysine treated grids. TbRRP44 knockdown cells, by their turn, presented a wide and flat conformation (supplementary Fig. S1). As described in the Materials and Methods section, a quick fixation step with 2% paraformaldehyde (PFA) blocked the flagellar movement of the wild-type cells and avoided deformation of the TbRRP44 knockdown cells, allowing the preparation of grids with cells evenly distributed without apparent deformations (supplementary Fig. S1).

An initial analysis of reconstructed 3D slices revealed that major organelles, including mitochondrion, nucleus, nucleolus, flagellar pocket, kinetoplast, lipid droplets, and other strong absorbing vacuoles showed regular conformation in control cells (Fig. 4). Consistently, 3D reconstruction of the cell structure and manual segmentation of the organelle surfaces shows control cells with nicely defined flagellum, kinetoplast, lipid droplets, a round flagellar pocket, a round nucleus, and the nucleolus. The mitochondrial network is well organized along the two sides of the cell (Fig. 4).

**Figure 4.**
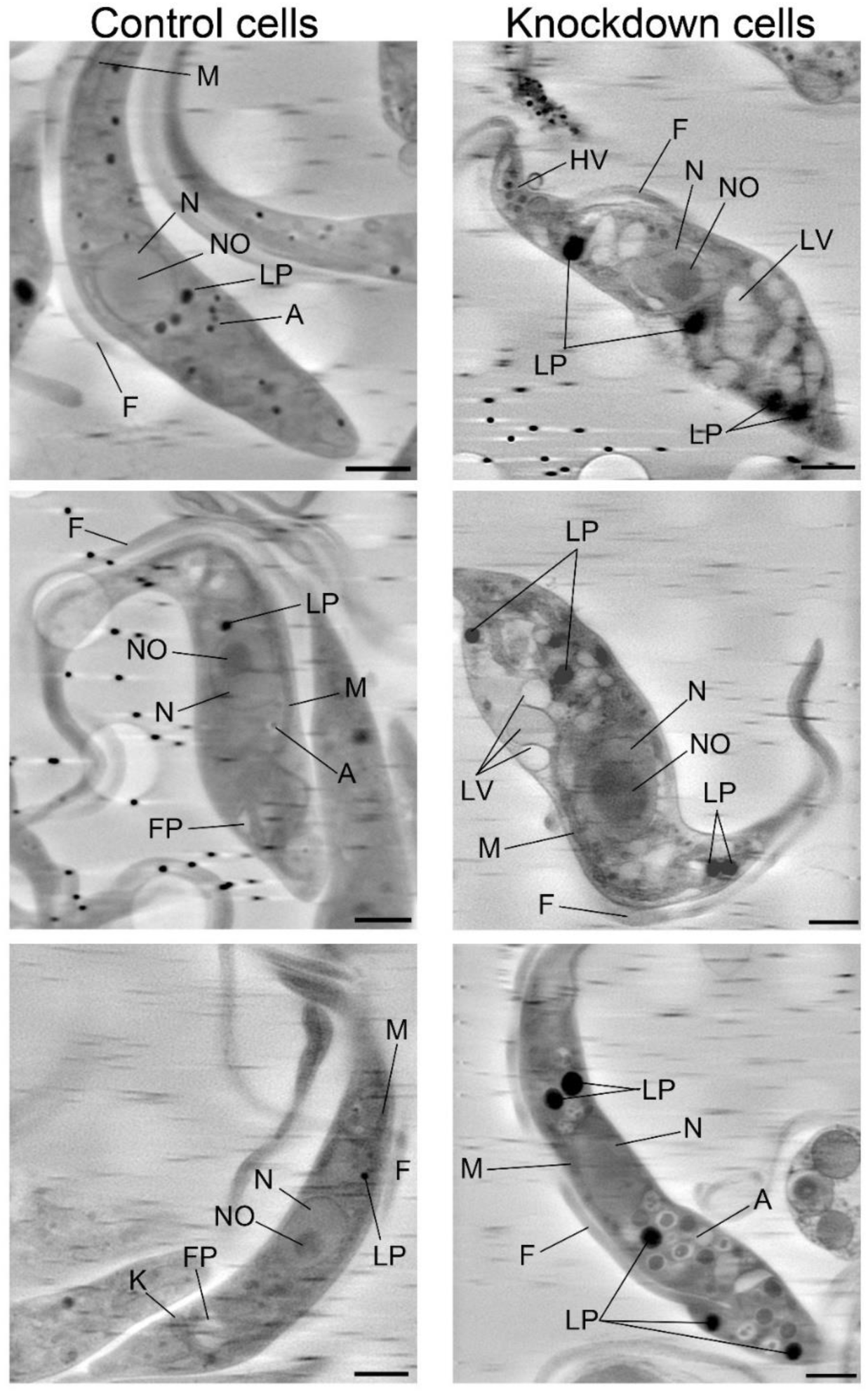
Cryo-soft X-ray tomography volume slices of control and TbRRP44 knockdown cells. Control cells are shown in the left panels. Knockdown cells at 72 hours after induction of RNA interference are shown in the right panel. Letters indicate the organelles identified. N: nucleus; NO: nucleolus; FP: flagellar pocket; F: flagellum; LP: lipid droplets; HV: high X-ray absorbing vesicles; A: acidocalcisomes; LV: low X-ray absorbing vacuoles; M: mitochondria. Scale bar, 1.5 µm.

TbRRP44 knockdown cells, on the other hand, presented structural changes with a certain degree of heterogeneity between different cells (Fig. 4). Some cells displayed a wider and shorter conformation, while others were elongated and thinner. However, a feature common to all knockdown cells was the extreme vacuolation of the cytoplasm and enlargement of all types of vesicles leading to a huge cytoplasmic crowding. The nuclei and nucleoli became larger, and the flagellar pocket appeared compressed (Fig. 4). Heterogeneities were observed both in the posterior and anterior poles of the cells. In the posterior pole, some cells showed low X-ray absorbing vacuoles and deformed kinetoplast, while others presented enlargement of vesicles with high absorbance (supplementary Fig. S2). Similarly, in the anterior pole, some cells contain a high number of large low absorbance vacuoles and others a high number of small vesicles with high X-ray absorption (supplementary Fig. S2). 3D reconstructions revealed different degrees of vacuolation in TbRRP44 knockdown cells (Fig. 5). The most striking alteration is the presence of a high number of low X-ray absorbing vacuoles that occupy most of the cytoplasmic space. Their absorption coefficient is within the range of the absorption coefficient presented by the flagellar pocket. As the low absorbing vacuoles take up most of the cytoplasm, they cause a displacement and disorganization of the mitochondrial network. Fig. 5 shows in the central panel a dividing cell with two kinetoplasts, a dividing flagellum, alterations in the mitochondrial network, and the presence of low absorbing vacuoles in both cell poles. The right panel shows a cell containing a single large vacuole taking most of the anterior pole and surrounding the nucleus. In most cells, the nuclear boundary is irregular, and the nucleolus is larger than in control cells. In addition, most TbRRP44 knockdown cells contain a higher number and larger lipid droplets (Fig. 5). Given the alterations revealed by cryo-soft X-ray tomography, we have performed additional analyses aiming at identifying the organelles affected by these alterations, which are described in the following sections.

**Figure 5.**
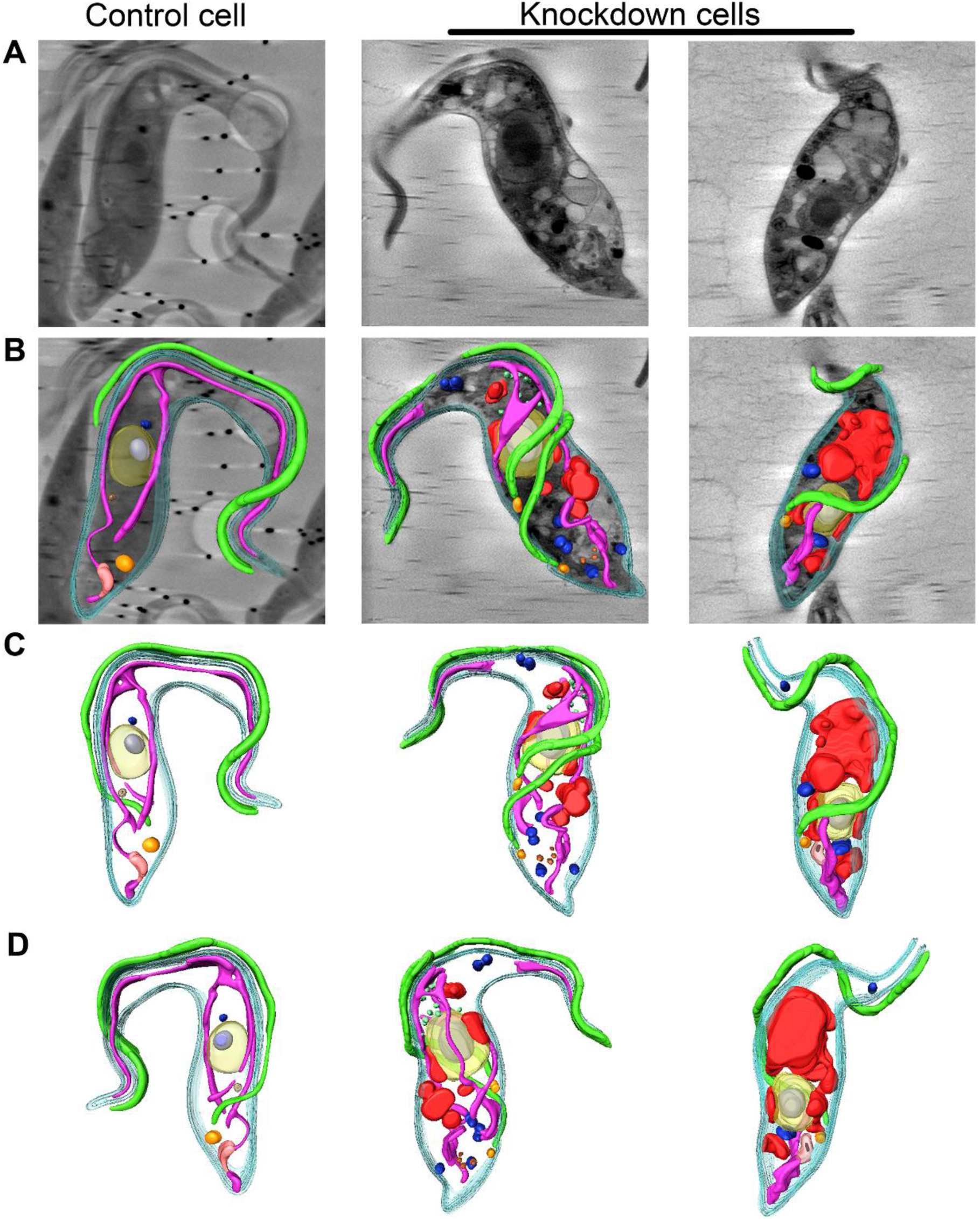
Segmentation of the 3D cryo-SXT reconstructions of *T. brucei* cells. A control cell is shown in the left panel. The center and right panels show cells after 72 hours of TbRRP44 depletion with different stages of vacuolation. The cell on the right shows a single large vacuole taking most part of the anterior pole. (**A**) Original volume slices before segmentation. (**B**) Organelle segmentation shown on a volume slice of control and knockdown cells. (**C-D**) Segmentation of cell structures with 180° rotation. Cyan: membrane; yellow: nucleus; light grey: nucleolus; green: flagellum; orange: flagellar pocket; purple: mitochondrion; pink: kinetoplast; brown: acidocalcisome; blue: lipid droplets; red: low X-ray absorbing vacuoles; light green: high X-ray absorbing vesicles.

### Acidocalcisomes present a larger size in TbRRP44 knockdown cells

Acidocalcisomes are lysosome-related organelles rich in calcium and other cations, also containing a high concentration of pyrophosphate (PPi) and polyphosphate (poly P). They possess proton pumps and transporters, playing a central role in calcium signaling and in phosphate and cation homeostasis in trypanosomatids ^60,61^. To evaluate the extent of alteration in calcium-containing organelles in *T. brucei* cells deficient for TbRRP44, we have used cryo spectromicroscopy at MISTRAL beamline to obtain X-ray absorption near-edge structure spectra (XANES) on fully hydrated cells, therefore avoiding Ca ion leakage. Sequential images were acquired from the same field of view starting below the Ca L2,3 absorption edge, up to 356 eV, which is above the Ca absorption edge ^62,63^ (Fig. 6A). For localization of calcium-containing organelles, images were then acquired at pre-edge and calcium L3-edge energies for several control and knockdown cells. Calcium was identified by the difference between the L3-edge and pre-edge images (Fig. 6C). We assumed that the organelles presenting high calcium content correspond to the acidocalcisomes. The average volume of acidocalcisomes is approximately 5 times larger in TbRRP44 knockdown cells when compared with control cells (Fig. 6B).

**Figure 6.**
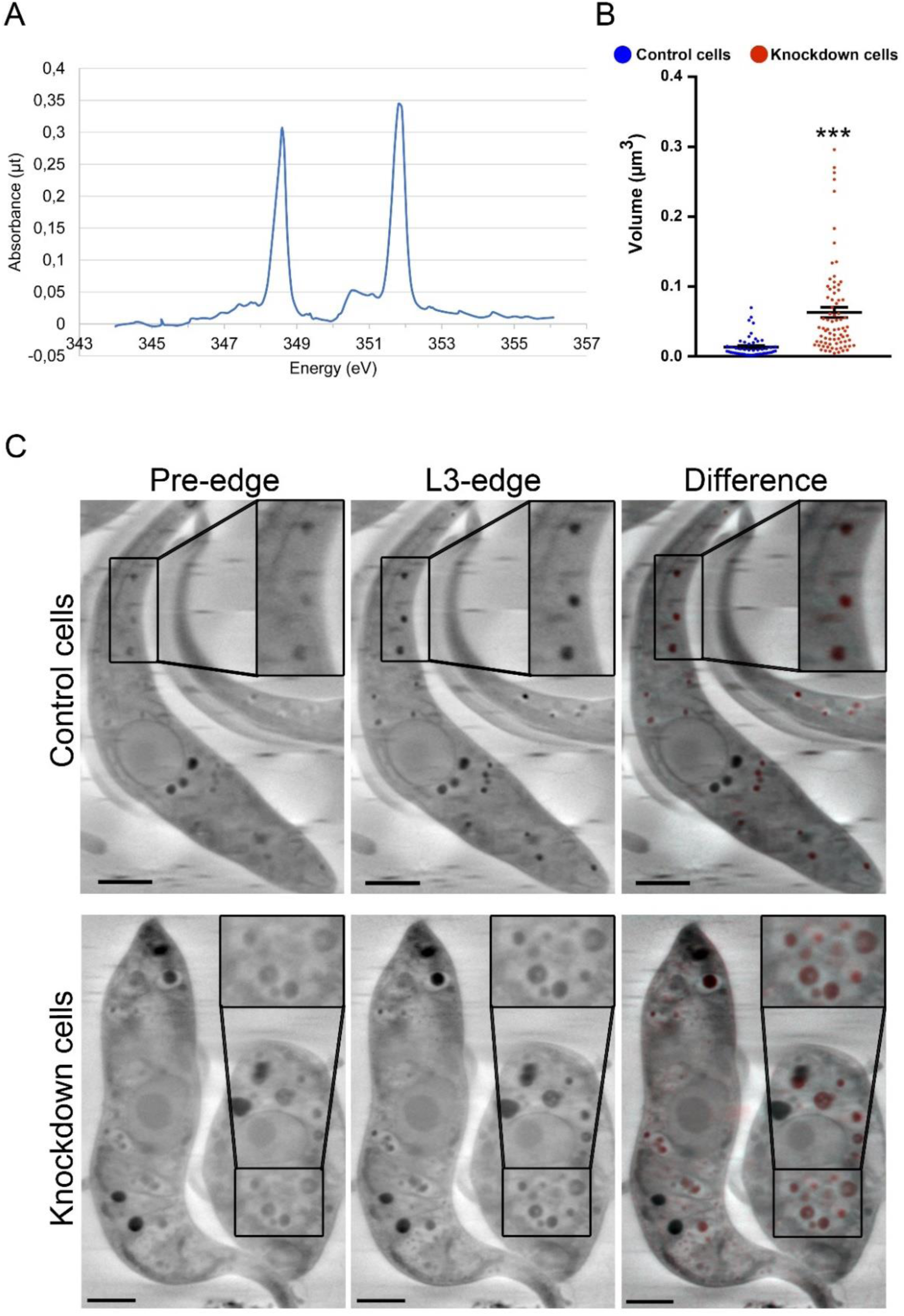
Cryo spectromicroscopy at the Ca L-edges. (**A**) XANES spectrum of Ca-containing vesicles showing the L3- and L2-edges (first and second peaks, respectively). The spectrum corresponds to Ca in solution as in ^63^. (**B**) Quantification of the volume of calcium-containing vesicles from three cryo-SXT reconstructed cells. (**C**) Representative images of control and knockdown cells collected at the calcium pre-edge (left panel) and L3-edge (center panel). The calcium-containing organelles evidenced by the difference between the L3-edge and pre-edge images are highlighted in red over an image of the same cell (right panel). Scale bar, 1.5 µm.

### TbRRP44 knockdown cells contain a higher number of lipid droplets showing a larger volume

The analyses with soft X-rays in the calcium pre- and absorption edges provided information that allowed us to distinguish the acidocalcisomes from the vesicles showing the strongest soft X-ray absorption (Fig. 6C). These vesicles correspond to lipid droplets, which display a higher absorption than water rich organelles ^64–67^. Lipids are essential components of cellular membranes and key components for energy storage and metabolism. Neutral lipids, such as fatty acids, are stored in lipid droplets, which are the major cellular reservoirs for this type of molecule. Lipid droplets also contain a specific set of proteins, such as enzymes involved in lipid synthesis, membrane trafficking, and organelle transport. They also protect against the toxic effects of free fatty acids excess [reviewed in ^68,69^]. Given the central role of these organelles for cell function and the interplay of fatty acids from lipid droplets with other organelles, we evaluated the size of lipid droplets in cells depleted of TbRRP44. They were quantified both from 3D cryo-SXT reconstructions and from images of cells stained with the fluorescent dye Nile Red acquired using a high-content imaging system, which allowed the analysis of a larger number of cells. The 3D-reconstructed knockdown cells showed an average of six lipid droplets per cell, while the average of control cells was approximately three (Fig. 7A and B). In addition, the lipid droplets in knockdown cells showed an average size two times larger than in control cells (Fig. 7B). In the analysis using Nile Red, TbRRP44 knockdown cells showed both a wider range in fluorescence signal intensity and a significant increase of fluorescence intensity average (Fig. 7C and D). However, the difference between control and knockdown cells is smaller relative to the quantifications of the lipid droplets by cryo-SXT 3D reconstructed cells. This fact could be due to the lower resolution of the fluorescence microscopy technique. Together, the quantifications showed that lipid droplets are highly affected in cells deficient for TbRRP44.

**Figure 7.**
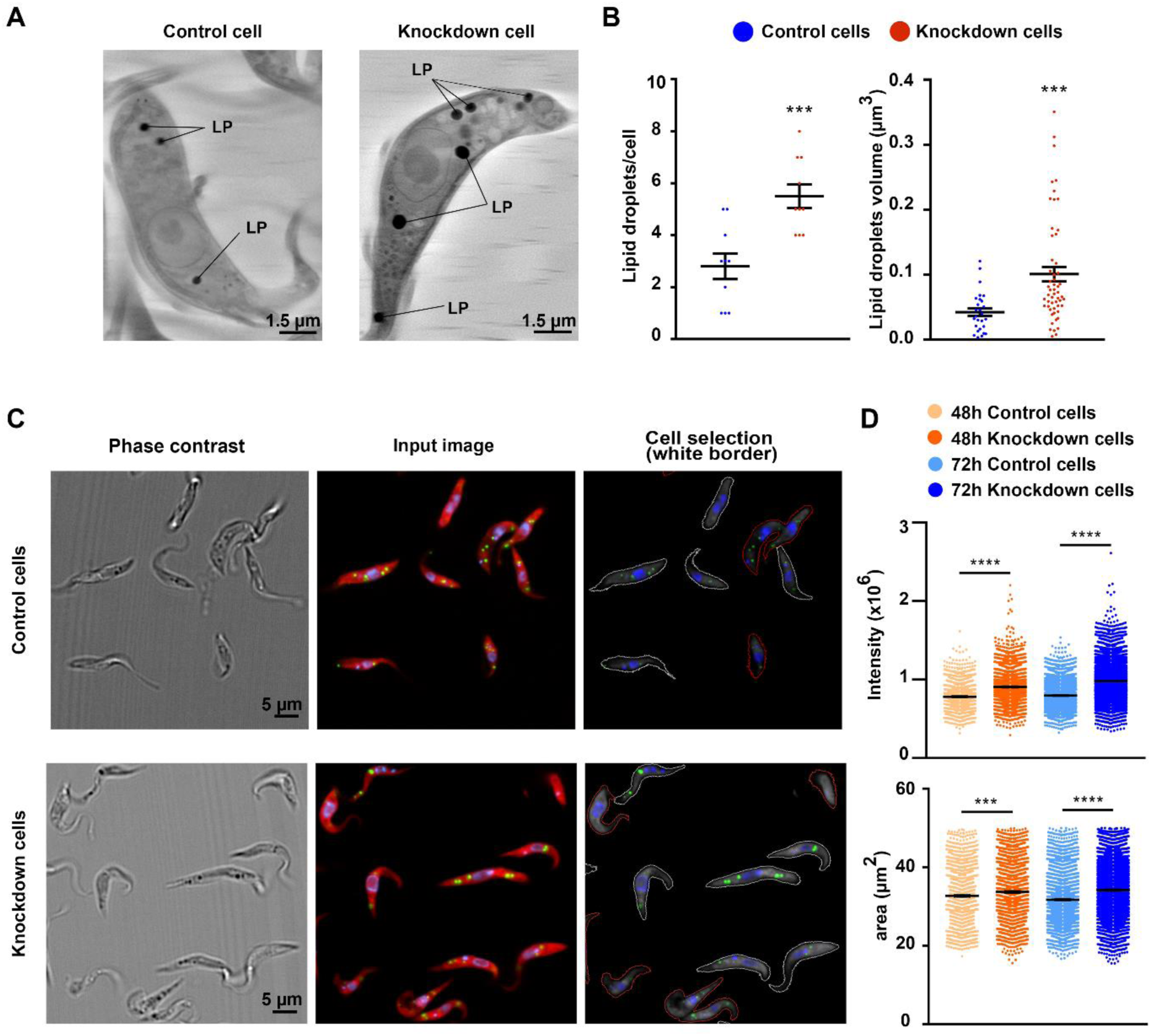
*T. brucei* TbRRP44-depleted cells present a higher number and larger lipid droplets. (**A**) Representative cryo-soft X-ray tomography volume slices of control (left) and TbRRP44 knockdown (right) cells. Lipid droplets are indicated (LP). (**B**) Number per cell (left) and volumes (right) of lipid droplet according to quantification from cryo-SXT reconstructed cells. A total of 10 cells was used in this analysis. (**C**) Representative images of the cells used for quantification of lipid droplets stained with Nile Red. Left: phase contrast images. Center: cells imaged in red, nuclei and kinetoplasts in blue and lipid droplets in green. Green and red fluorescence are both from Nile Red. Right: Cell contour for quantification of cell area defined based on the red fluorescence. (**D**) Quantification of the green fluorescence intensity and area of cells stained with Nile Red. Knockdown cells were collected at 48 and 72 hours after RNAi induction.

### Mitochondrial activity is affected TbRRP44 knockdown cells

As described above, 3D reconstructions revealed disorganization of the mitochondrial structure in TbRRP44 knockdown cells. To evaluate the state of the mitochondrion in these cells, we have used the fluorescent dye MitoTracker, which accumulates in the mitochondria of living cells once added to cultures at low nanomolar concentrations. In active mitochondria, it undergoes oxidation and emits fluorescence (Fig. 8A). The fluorescence intensity average per cell area was two times lower in TbRRP44 knockdown cells when compared to control cells (p < 0.001) (Fig. 8B). This decrease indicates a reduction or loss of mitochondrial membrane potential affecting mitochondrial activity. This analysis also confirmed that TbRRP44 knockdown cells show an increase in the cell area (control cells, 31.2 µm^2^; knockdown cells, 37.6 µm^2^; p < 0.001) (Fig. 8B).

**Figure 8.**
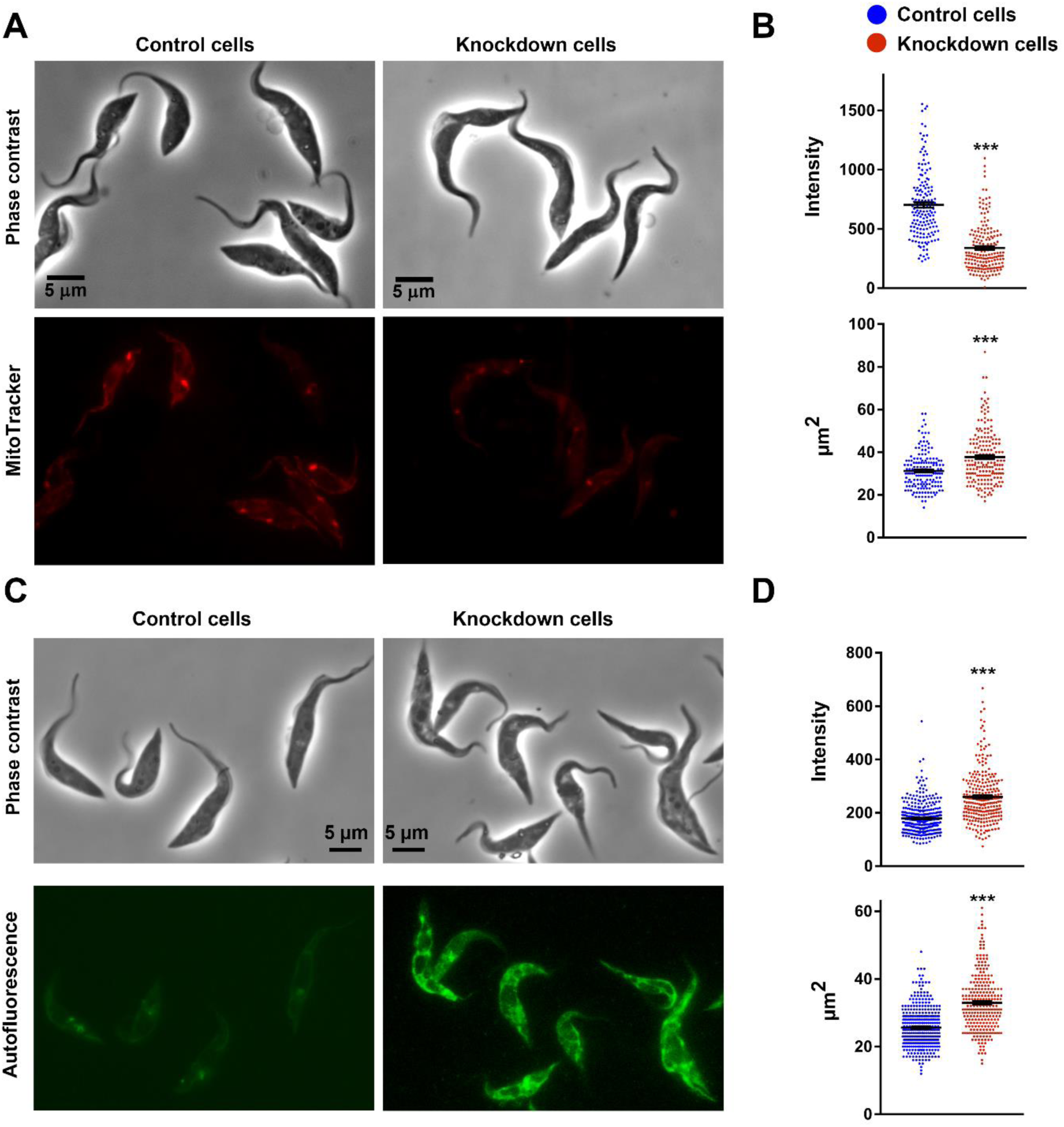
TbRRP44 knockdown affects mitochondrial activity. (**A**) Representative images used in the analysis of mitochondrial activity as determined by MitoTracker oxidation. Upper panel: phase contrast images. Lower panel: fluorescence images of cells stained with 10 nM MitoTracker Orange CMTMROS. Left: control cells. Right: TbRRP44 knockdown cells. (**B**) Quantification of MitoTracker fluorescence intensity (upper graph) and cell area (lower graph). (**C**) Representative images used in the analysis of *T. brucei* autofluorescence in the green wavelength range. Upper panel: phase contrast images. Lower panel: fluorescence images with excitation at 460-500 nm and emission at 512-542 nm. Left: control cells. Right: TbRRP44 knockdown cells. (**D**) Quantification of fluorescence intensity (upper graph) and cell area (lower graph).

It is known from a previous study that cells from the trypanosomatid *Leishmania tarantolae* display significant autofluorescence with excitation at 450-500 nm and emission peaks at 458/538 nm ^70^. The major contributor to fluorescence in this range is Flavin adenine dinucleotide (FAD)^71^. FAD and its reduced form FADH2 are involved in catabolic reactions, electron transport chain, and energy metabolism, but only FAD presents autofluorescence. In addition, FAD exists mostly as a cofactor bound to enzymes involved in redox reactions ^71,72^. Autofluorescence in the FAD contribution range was determined for *T. brucei* control and cells depleted of TbRRP44 for 72 hours. As expected, *T. brucei* also showed a relatively strong autofluorescence. In control cells, autofluorescence appears to colocalize with the single giant mitochondrion ^73^, being distributed along the cell membrane with a dark central area, which is occupied by the nucleus and adjacent structures. Surprisingly, TbRRP44 knockdown cells presentedled to a stronger autofluorescence, with an average approximately 50% higher than control cells (Fig. 8C and D). These cells also presented a larger area average (32.9 µm^2^) relative to the controls (25.6 µm^2^) (p < 0.001) (Fig. 8D). Considering that fluorescence in this range is attributed to the oxidized FAD form, this increase suggests that energy metabolism is affected in TbRRP44 knockdown cells and is consistent with the results obtained with MitoTracker staining, which indicated alterations of mitochondrial activity in TbRRP44-depleted cells.

### Acidic and lysosome-derived vacuoles show a high increase in size and number in TbRRP44 knockdown cells

In order to investigate the relation of the vesicles observed in TbRRP44-depleted with the lysosome, we performed analyses using the chemical marker LysoTracker and the lysosomal/endosomal protein p67 marker. Despite its name, LysoTracker can be incorporated into all acidic vesicles in living cells, which in the case of *T. brucei* should include lysosomes and acidocalcisomes. Interestingly, TbRRP44-depleted cells showed very strong fluorescence upon staining with LysoTracker. The ratio of total fluorescence intensity per cell area revealed a six-fold difference between control and knockdown cells (Fig. 9A and B). In addition, the LysoTracker-stained vacuoles are much larger in TbRRP44 knockdown cells. These findings are consistent with the hypothesis that the vacuoles which increase in size and number originate from lysosomes.

**Figure 9.**
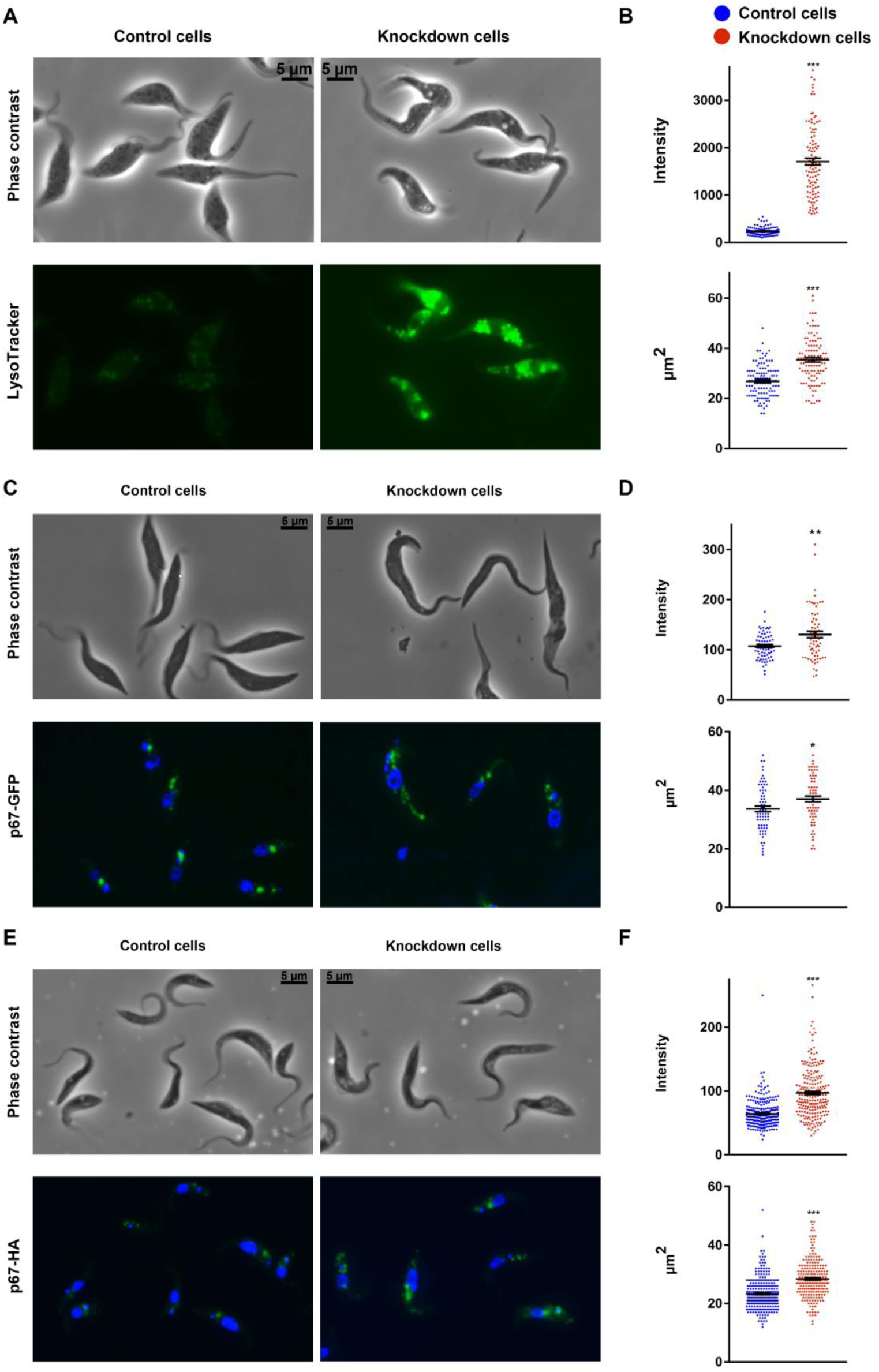
Increase of acidic and lysosome-derived vacuoles in TbRRP44 knockdown cells. (**A**) Representative images used in the analysis of acidic vacuoles as determined by LysoTracker labeling. Phase contrast and LysoTracker fluorescence images are shown in the upper and lower panel respectively. Control cells and TbRRP44 knockdown cells are shown in the left and right panels. (**B**) Quantification of LysoTracker fluorescence intensity (upper graph) and cell area (lower graph). (**C**) Representative images used in the analysis of the lysosomal protein marker p67 localization using p67 tagged with GFP. (**D**) Quantification of fluorescence intensity (upper graph) and cell area (lower graph) of cells expressing p67-GFP (**E**) Representative images used in the immunolocalization analysis of the lysosomal protein p67 tagged with HA. (**F**) Quantification of fluorescence intensity (upper graph) and cell area (lower graph) of cells expressing p67-HA immunostained with anti-HA antibody.

This hypothesis was further supported by the analysis of the lysosomal/endosomal membrane protein p67 localization. This protein was previously shown to localize in *T. brucei* lysosomes in fusion with both the yellow fluorescent protein (YFP) ^74^ (Fig. 9C and D) and the influenza hemagglutinin (HA) epitope tag ^75^ (Fig. 9E and F). We have made equivalent fusions to p67 and evaluated its subcellular localization in the TbRRP44 conditional strain. Fluorescence microscopy analysis revealed a higher number of fluorescent vacuoles for p67 fused both to the HA tag and the green fluorescence protein. Total p67-GFP fluorescence was significantly increased in TbRRP44 knockdown cells. However, as the knockdown cells present a larger area average (control cells, 33.7 µm^2^; knockdown cells, 37 µm^2^; p = 0.016), the difference in GFP fluorescence intensity per cell area between control and knockdown cells was not significant (Fig. 9C and D). Immunolocalization of p67-HA using an anti-HA antibody, on the other hand, revealed a significant increase of p67-positive vacuoles in the quantification of total fluorescence, in the ratio between the fluorescence intensity, and in the cell area, even though TbRRP44 knockdown cells presented a larger size (control cells, 23 µm^2^; knockdown cells, 28 µm^2^; p < 0.001) (Fig. 9E and F). These results are consistent with the hypothesis that the large vesicles should be derived from lysosomes. We can conclude that the low X-ray absorbing vacuoles correspond to the vesicles stained with both lysotracker lysosomal marker p67.

## Discussion

The evidence available so far indicates that the biochemical activity of Rrp44/Dis3 homologues is required for accurate processing and maturation of many pre-RNA types as well as for maintenance of posttranscriptional RNA homeostasis. While the absence of Rrp44/Dis3 activity is lethal, a wide range of phenotypes have been reported associated with partial Rrp44/Dis3 deficiency, affecting chromosome segregation, mitotic progression, genome stability, and function ^1,2,25,27–30,37^, and, embryo development in multicellular organisms ^22–26^. In these organisms, its function is also required for the control of cell differentiation and proliferation in some specific tissues ^25,31,32,39–41^.

Based on the different biological outcomes caused by Rrp44/Dis3 deficiency in other organisms, we can expect that TbRRP44 deficiency may also lead to phenotypes particular to *T. brucei* cells. The ratio of autophagosomes per cell in TbRRP44-depleted cells is consistent with previous reports on autophagy induction in *T. brucei* ^51,76–78^ and supports the hypothesis that TbRRP44 depletion activates the autophagy response. Autophagy has been linked to cell death in *T. brucei* ^49^ and correlates with the mammalian cell response to nucleolar stress caused by deficient ribosome synthesis, which triggers its ^47,48^. Therefore, autophagy may represent one of the components that sends inhibitory signals to other intracellular organelles in TbRRP44-depleted cells.

Our data show that, while the intracellular organelles start to degrade, the cell membrane and the nuclear DNA continue to resist longer, showing signs of instability and degradation only by late times of TbRRP44 depletion. This relatively low level damage indicates that TbRRP44 knockdown cells are not following an apoptosis-like cell death process, as the one described for spliced leader silencing ^79,80^ and knockdown of the nuclear RNA-binding protein TbRRM1 ^54^. In these cases, the cell death process follows with a strong increase in the hypodiploid sub-G1 population due to nuclear DNA and cell membrane damages. Trypanosomatids do not seem to have the same molecular machinery that regulates cell death by apoptosis in multicellular organisms. It has also been proposed that trypanosomatid cell death occurs either by necrosis or by incidental death [reviewed in ^81^]. TbRRP44 knockdown cells present increased size, possibly related to oncosis, in parallel with translucent cytoplasm and swelling of organelles, which have been associated with necrosis ^81,82^. These features suggest that different cellular processes converge to induce cell death after the knockdown cells stop proliferating.

Since depletion of TbRRP44 does not lead to an increase in DNA content, the finding that these cells contained nuclei with larger size was intriguing. Similar events have been reported for mammalian cells undergoing senescence ^55–57^. To our knowledge, such a process has not been described for trypanosomatids yet, but it is well known for the budding yeast *S. cerevisiae*, which undergoes a limited number of divisions before entering senescence and dying. During the senescence process, *S. cerevisiae* has been recently shown to accumulate extra chromosomal rDNA circles that lead to a massive increase of unprocessed pre-rRNA in the nucleolus, resulting in loss of nuclear homeostasis with the increase of the nucleus size and accumulation of nuclear proteins ^83^. TbRRP44 knockdown cells and senescent *S. cerevisiae* cells do share one feature, which is the accumulation of unprocessed pre-rRNA. However, currently, we lack for *T. brucei* the same set of genetic and immunological tools that are available for *S. cerevisiae* and mammalian cells to experimentally confirm that accumulation of unprocessed pre-rRNA affects nucleolar and nuclear homeostasis and participates in the signals that lead to inhibition of cell proliferation in *T. brucei*.

The analyses performed by cryo-soft X-ray tomography provided detailed evidence of the ultrastructural alterations that take place in TbRRP44 knockdown cells (Fig. 4 to 7). The alterations involve the nucleus, nucleolus, and mitochondria localization and structure, along with a general enlargement of all vesicles. The most striking alteration involves formation of low X-ray absorbing vacuoles. Reconstitution of the three-dimensional cell structures showed that these vacuoles can increase in size and number and occupy most of the cytoplasmic space. The vesicles stained with LysoTracker are the ones showing the highest increase in size in TbRRP44 knockdown cells. A similar pattern was observed for the vesicles detected by the lysosomal membrane protein p67 marker. Therefore, the large acidic vesicles should be derived from lysosomes and correspond to the low X-ray absorbing vacuoles observed in the 3D cryo-soft X-ray reconstructions. The formation and expansion of lysosome-derived structures correlate with cell death processes in *T. brucei* ^43–46^. The different degrees of vacuolation suggest that TbRRP44 knockdown cells are at different stages of a death process, with the cytoplasm being taken up by lysosome-derived vacuoles.

Acidocalcisomes were localized by elemental specific imaging at the calcium pre- and L2,3-edges. Volume calculation confirmed that acidocalcisomes increase in size in TbRRP44 knockdown cells, but they are not the vesicles that make the largest contribution to vacuolation. Our analyses also provided direct evidence to distinguish the acidocalcisomes from lipid droplets, which contain lipids at high density, and are the organelles showing the highest absorption in the “water window” energy range ^64–66^. Both the number per cell and the size of lipid droplets observed in the 3D reconstructed knockdown cells are two times higher than in control cells. Nile Red fluorescence was useful to quantify lipid droplets, although at a lower resolution than cryo-SXT. The increase in size and number of lipid droplets contributes to the general enlargement of vesicles in TbRRP44 knockdown cells and indicates an unusual accumulation of lipids in these cells.

Despite presenting a disorganized structure in the cryo-SXT 3D reconstructions, the analyses with the mitochondrial marker MitoTracker did not indicate that the mitochondrion is a major contributor to the general enlargement of vesicles in TbRRP44 knockdown cells. However, these cells showed loss or reduced mitochondrial membrane potential as detected by their failure to oxidize MitoTracker (Fig. 8). In addition, the strong increase of autofluorescence in the green range (512-542 nm), where FAD is the major contributor and the mitochondrion identified as its major source ^70^, is consistent with strong alterations of energy metabolism in these cells. Loss of the mitochondrial membrane potential is also one of the initial signals of activation of cell death mechanisms.

In conclusion, the multidisciplinary approach used in this work provided for the first time a detailed description of drastic ultrastructural cellular defects that are taking place in *T. brucei* cells, after depletion of an essential protein, the TbRRP44 ribonuclease. Our results revealed extensive cell alterations, including phenotypes never described before, which can add to the knowledge of cell death mechanisms in trypanosomatids.

## Materials and methods

### *Trypanosoma brucei* 29-13 derivative strains

Construction of a *Trypanosoma brucei brucei* [here designated as *Trypanosoma brucei* (*T. brucei*) for simplification] strain for conditional depletion of the TbRRP44 was described in a previous work of our group ^42^. Procyclic cells of the parental *T. brucei* 29–13 strain were maintained in SDM 79 supplemented with 10% (v/v) bovine calf serum, hygromycin (50 µg/mL) and G418 (15 µg/mL). The medium for the conditional strain also contained phleomycin (2.5 µg/mL). The conditional strain was also transformed with plasmid pGL2166 to express the autophagy marker Atg8.2 fused to the yellow fluorescence protein. Plasmid pGL2166 ^51^ was kindly provided by Jeremy Mottram (Centre for Immunology and Infection, University of York, UK).

### Induction of RNA interference and cell fixation with paraformaldehyde (PFA)

Depletion of TbRRP44 was performed as described previoulsy^20^.Briefly, cells from fresh cultures at a density of 1-5x10^7^ cells/mL were inoculated in two cultures of 10 mL SDM 79 supplemented with 10% (v/v) bovine calf serum containing hygromycin (50 µg/mL), G418 (15 µg/mL) phleomycin (2.5 µg/mL) and maintained at 28°C. RNA interference was induced by adding tetracycline (2 µg/mL at time 0 followed by 1 µg/mL every 24 hours). Cells from the TbRRP44 conditional strain were collected at 48, 72, and 96 hours after RNAi induction. For each time point, samples grown in parallel without induction with tetracycline were collected as controls. After each incubation time, cells were centrifuged at 3000 x g for 5 min, washed with phosphate buffered saline (PBS, 10 mM sodium phosphate buffer pH 7.4, 138 mM NaCl, 27 mM KCl), and fixed with 4% (v/v) paraformaldehyde (PFA) in PBS at room temperature for 10 min. Subsequently, the cells were washed again, suspended in PBS, and maintained at 4°C for analysis.

### Amino acid starvation and autophagosome analyses

Autophagosomes were analyzed using the TbRRP44 conditional strain transformed with plasmid pGL2166 expressing the autophagosome marker ATG8.2 fused to the yellow fluorescent protein ^51^. The cells were cultivated under permissive and knockdown conditions and collected for analysis at 48 and 72 hours after the RNAi induction. Induction of autophagosome formation was carried out as described by Li et al. ^49^ using gPBS (PBS containing 1 g/L glucose) instead of gHBSS. The cells were washed twice with gPBS (PBS containing 1 g/L glucose), suspended in gPBS at 1-3 x 10^6^ cells/mL, and incubated at 28°C for 2 hours. Subsequently, the cells were fixed with 4% PFA as described above and adhered to poly-L-lysine-treated microscope slides. The cells were permeabilized with 0.25 % (v/v) NP40 in PBS buffer for 2 min and blocked with 4% bovine serum albumin in PBS buffer for 1 hour. Atg8.2-YFP was localized by immunostaining with an anti-GFP nanobody [clone LAg16-G4S-2 ^84^] conjugated with Alexa Fluor 488 and simultaneously stained with a 1:1500 dilution of a 1% Evans Blue (v/v) solution. Images were acquired on a Leica Microsystems DMI6000B microscope using an HCX PL APO 100x/1.40 oil objective. This microscope is equipped with a DFC365 FX camera and operates with Leica Application Suite. Acquisition time was 0.5 seconds (460-500 nm bandpass filter for excitation and a 512-542 nm bandpass filter for emission) and 0.1 seconds for phase contrast images. The images were analyzed with Columbus Image Data Storage and Analysis System for Microscopy (PerkinElmer). Cells were detected using the “find cell” algorithm (method A) from the Evans blue signal (red channel). Objects near borders were excluded, and only entire cells were included in the analysis. Isolate cells were selected by morphology (roundness and area) properties. Autophagosomes were identified using the Alexa fluor 488 signal (green channel) with the “find spots” algorithm (method C) and the percentage of cells with autophagosomes was calculated. The data were obtained from 10 images of each experimental condition. The total number of cells analyzed in each experimental conditions was as follows: control cells at 48 hours, 514 cells incubated in SDM79 and 743 cells incubated in gPBS; TbRRP44-knockdown cells at 48 hours, 524 cells incubated in SDM79 and 439 cells incubated in gPBS; control cells at 72 hours, 397 cells incubated in SDM79 and 328 cells incubated in gPBS; TbRRP44-knockdown cells at 72 hours, 312 cells incubated in SDM79 and 331cells incubated in gPBS.

### Cell cycle analysis by flow cytometry

For cell cycle analyses, control and TbRRP44 knockdown cells were collected 48, 72 and 96 hours after RNAi induction. The cells were centrifuged at 300 x g for 10 min, washed with PBS, and fixed with cold ethanol 70% (5 x 10^6^/mL) at -20 °C. After that, the cells were harvested by centrifugation, washed with PBS, and suspended in 200 µL PBS plus 200 µL of a propidium iodide staining solution (3.4 mM Tris.HCl pH 7.4, 0.1% NP40 (v/v), 10 mg/mL RNase A, 10 mM NaCl, 30 µg/mL propidium iodide). Data from 10,000 events per sample were acquired using a FACS Canto II (BD Biosciences). The propidium iodide fluorochrome was excited with a blue laser (488 nm) and the emitted fluorescence was collected through a 585/42 bandpass filter. Single cells were gated based on PI-A vs. PI-W ^85^. The cell samples were stained and analyzed in triplicate. Data analysis was performed using the FlowJo software version 10.6.1.

### Analysis of cell death signals using annexin V-Alexa 488 and propidium iodide staining

A total of 2x10^6^ cells from control and TbRRP44 knockdown cells from 48, 72, and 96 hours after RNAi induction were centrifuged at 300 x g for 10 min, washed once with PBS, and stained with Alexa Fluor 488 annexin V/Dead Cell Apoptosis Kit (Invitrogen - V13245) according to the manufacturer’s instructions. The analyses (20,000 events/sample) were performed on a FACS Canto II (BD Biosciences). Alexa Fluor 488-annexin V and propidium iodide fluorochromes were excited with a blue laser (488 nm), and the emitted light was collected through 530/30 and 585/42 band pass filters, respectively. Color compensation was used to minimize spectral overlap. The cell samples were stained and analyzed in triplicate. Data analysis was performed using the FlowJo software version 10.6.1.

### Nucleus and kinetoplast quantification

Control and TbRRP44 knockdown cells were collected at 48 and 72 hours after RNAi induction and fixed with 4% (v/v) PFA (∼3 x 10^6^ cells/mL) as described above. The cells were adhered to CellCarrier-96 Ultra Microplates (PerkinElmer) precoated with poly-L-lysine and stained with 2 µg/mL of DAPI (4’,6-Diamidino-2-Phenylindole). The images were acquired on an Operetta CLS High-Content Analysis System (PerkinElmer) using the confocal model, z-stack function (four planes at intervals of 0.5 µm) and a 63x water objective lens (NA 1.15 and WD 0.6). The DAPI signal was collected using 355-385 nm excitation and 430-500 nm emission filters. Maximum projection images from Z-planes were analyzed with Harmony 4.6 software. Cells were detected using the “find cell” algorithm (method A) in the digital phase contrast (high detail) images. Cells close borders were excluded, and only entire cells were kept in the analysis. Isolate cells were selected based on morphological properties (width and area). Nuclei and kinetoplasts were identified as spots (method B) in DAPI images and were distinguished based on the area and roundness properties. The output results represent the percentage of 1N/1K, 1N/2K, and 2N/2K cells.

### Cell preparation for cryo-soft X-ray tomography and cryo spectromicroscopy

Induction of RNA interference was performed as described above, and cells were collected at 48 and 72 hours after induction. In initial attempts of grid preparation, TbRRP44 knockdown cells appeared to be very fragile as they “compressed”, assuming a flat conformation (supplementary Fig. S1). On the other hand, wild type procyclic *T. brucei* cells are fast moving and did not adhere well to poly-L-lysine treated grids. Both problems were overcome by a quick fixation step with 2% paraformaldehyde (PFA) as follows: 1 mL of a cell culture containing ∼10^7^ cells/mL was centrifuged, washed in 500 μL PBS, quick-fixed in 500 μL PBS containing 2% PFA (v/v) for 3 min at room temperature, and washed again with 500 μL PBS. All centrifugation steps were at 800 x g for 5 min at room temperature. The PFA quick-fixed cells were suspended 100 μL PBS at ∼10^8^ cells/mL. Before cell deposition, the carbon side of gold quantifoil R 2/2 holey carbon-film microscopy grids (Au-G200F1) were treated with 20 μL of poly-L-lysine in a humidity-controlled chamber at 37°C for 30 min. The grids were washed twice with MillliQ water, and 4 μL of 100 nm gold nanoparticles (5.6 x 10^9^/mL) were deposited on each grid, which were let dry under a clean bench. 5 μL of cells (∼10^8^ cells/mL) were deposited on the grids and vitrified by plunge freezing using a Leica Automatic Plunge Freezer EM GP2. The PFA treatment preserved the morphology of the cells and organelles, which allowed for comparison of the volumes and ultrastructural alterations that occur upon downregulation of TbRRP44. However, PFA interferes with the linear absorbance coefficient (LAC) (supplementary Fig. S1), preventing quantitative comparison of LAC between the organelles of control and TbRRP44 knockdown cells, as well as comparison with the LAC of other tissues.

### Acquisition of cryo-soft X-ray tomography data sets and 3D volume reconstruction

Cryo soft X-ray tomography data were collected at the MISTRAL beamline (Synchrotron ALBA, Barcelona, Spain) ^86^. The vitrified grids were transferred to the beamline under cryogenic and vacuum conditions. The data sets were acquired at 520 eV energy with 1 second of exposure, in a tilt range from -70 to 70 degrees, rotating 1 degree between each image. Alignment of the tilted series was performed with IMOD ^87^ using the gold fiducial markers. Final reconstructions were performed using SIRT in Tomo3D ^88^, and cell segmentation was performed using the Amira 3D Software for Life Sciences (ThermoFisher Scientific).

### Cryo-soft X-ray spectromicroscopy

In order to analyze the calcium content in the cells, cryo spectromicroscopy was performed, and XANES spectra of the Ca-containing vesicles were analyzed by plotting the average intensity of the pixels of the vesicles area as a function of the energy (from calcium pre-edge to calcium post-edge). All vesicle spectra showed the Ca in solution form ^58^. Calcium localization in different control and TbRRP44 knockdown cells was carried out by subtracting a projection image of the cell taken at pre-edge from the one taken at the L3-edge. The contrast in the difference projection is dominated by areas containing substantial Ca content. Projection subtractions were carried out using the Fiji software ^89^. The volumes of acidocalcisomes were calculated using the Microscopy Image Browser software platform ^90^. The organelles were manually segmented, the volumes obtained in voxels (pixel^3^), and then converted into µm^3^. Statistical analyses were performed using GraphPad Prism version 7.01 software.

### Lipid droplet quantification

The volumes of lipid droplets from cryo-SXT reconstructed cells were calculated as described above for the acidocalcisomes. For Red Nile staining of lipid droplets, 2 mL of control and TbRRP44 knockdown cells were collected 48 and 72 hours after RNAi induction, fixed with 4% PFA (∼3 x 10^6^ cells/mL) as described above and adhered to 96-well plate pre-coated with poly-L-lysine. The cells were incubated with 1µg/mL of Nile red and 2 μg/mL of DAPI (4’,6-Diamidino-2-Phenylindole) for 30 min. The images were acquired on an Operetta CLS High-Content Analysis System (PerkinElmer) using the confocal model, z-stack function (three planes at intervals of 0.5 µm) and a 63x water objective lens (NA 1.15 and WD 0.6). The DAPI signal was collected using 355-385/430-500 nm excitation/emission filters. Nile red signals were collected using 460-490/500-550 (green) and 615-645/655-760 (red) excitation/emission filters. Maximum projection images from Z-planes were analyzed with Harmony 4.6 software. Cells were detected using the “find cell” algorithm (method C) using the red images. Border cells were excluded, and only entire cells were kept in the analysis. Isolated cells were selected based on morphology properties (width, length, area, and roundness). The intensity average of the green fluorescence signal from the selected cells was used for comparisons.

### PCR-based tagging of the lysosomal protein marker p67

The G418 genetic resistance marker from plasmids pMOTAG3H and pMOTAG3G ^91^ was replaced by the puromycin resistance gene. The resulting plasmids (pMO-HA-Puro and pMO-GFP-Puro) were used in PCR reactions to amplify the puromycin gene flanked by sequences of the target genes for introducing C-terminal tags into the lysosomal/endosomal membrane protein p67 (p67.1, gene ID: Tb427.05.1810). The PCR products were transformed into the *T. brucei* strain for conditional depletion of TbRRP44 ^42^. Following transfection, homologous recombination of the PCR products results in insertion of the tags sequences in the respective genes together with the puromycin resistance gene, which was used for selection of positive transfectants. Following selection with puromycin (2 μg/mL), transfectant cells were maintained in SDM 79 supplemented with 10% (v/v) bovine calf serum, hygromycin (50 μg/mL), G418 (15 μg/mL), phleomycin (2.5 μg/mL) and puromycin (4 μg/mL). Double-stranded RNA expression for RNA interference was induced with tetracycline (2 μg/mL at time 0 followed by 1 μg/mL every 24 hours), as described above.

### Subcellular localization of the lysosomal protein marker p67

For analysis of p67-GFP localization, live procyclic control and TbRRP44 knockdown cells were collected 48 and 72 hours after RNAi induction, concentrated to ∼10^8^ cells/mL, and mounted on microscope slides. Images were acquired on a Leica Microsystems DMI6000B microscope using the same objective and set of filters described above for autophagosome analyses except for the acquisition time that was 1 second and 1.5 seconds for the cells collected at the 48- and 72-hour time points, respectively. For indirect immunofluorescence analysis of HA-tagged p67, control and TbRRP44 knockdown cells were collected at 48 and 72 hours after RNAi induction, fixed with 4% (v/v) PFA (∼3 x 10^6^ cells/mL) as described above, and adhered to slides precoated with poly-L-lysine. To immunolabel p67-HA, the cells were permeabilized and blocked with saponin solution (1% v/v saponin, 2% w/v bovine serum albumin in PBS buffer) for 1 hour. Subsequently, the cells were incubated with the anti-HA high affinity antibody (3F10) (1:500 dilution, Roche – cat. 11867423001), followed by incubation with an anti-rat secondary antibody conjugated to Alexa Fluor 488 (1:2000 dilution, Molecular Probes - cat. A21210), and 2 μg/mL of DAPI. Images were acquired on a Leica Microsystems DMI6000B microscope using the same objective and set of filters described above. The acquisition time of p67-HA fluorescence images was 0.5 seconds using a 460-500 nm bandpass filter for excitation and a 512-542 nm bandpass filter for emission. The images were rescaled using the Analyze Menu tools of Fiji (http://imagej.net/Fiji; https://imagej.nih.gov/ij/) ^92^ to convert image size from pixels to μm. Definition of regions of interest, subtraction of background, and fluorescence and area quantitation were performed using Fiji ^92^.

### Autofluorescence in the green range and staining with Lysotracker and MitoTracker

For analysis of autofluorescence in the green wavelength range, control and TbRRP44 knockdown cells were collected 72 hours after RNAi induction and fixed with 4% PFA (v/v) as described above. Autofluorescence in the green wavelength range was acquired on a Leica Microsystems DMI6000B microscope using an HCX PL APO 100x/1.40 oil objective. This microscope is equipped with a DFC365 FX camera and operated with Leica Application Suite. Acquisition time was 5 seconds for autofluorescence (460-500 nm bandpass filter for excitation and a 512-542 nm bandpass filter for emission) and 0.1 seconds for phase contrast images. LysoTracker staining was performed on living cells. Control and TbRRP44 knockdown cells were collected 72 hours after RNAi induction, washed with PBS, stained with 10 nM of LysoTracker Green DND-26 (Molecular Probes/ThermoFisher), and mounted on microscope slides. The images were acquired on a Leica Microsystems DMI6000B microscope using the same objective and set of filters as described above for autofluorescence, using 0.5 seconds of acquisition time. For MitoTracker staining, 10 nM MitoTracker Orange CMTMROS (Molecular Probes/ThermoFisher) were added to cultures of control and TbRRP44 knockdown cells 72 hours after RNAi induction. The cells were incubated for further 30 minutes, washed with PBS, and fixed with 4% (v/v) PFA as described above. Images of MitoTracker-stained cells were acquired on a Nikon Eclipse 80i microscope using a Planfluor 100x/1.30 objective, Evolution MP Color camera (Media Cybernetics) and QCapture Pro 6.0 software. The acquisition time of MitoTracker fluorescence images was 4 seconds using a 528-553 nm bandpass filter for excitation and a 590-650 nm filter for emission. The images were rescaled using the Analyze Menu tools of Fiji (http://imagej.net/Fiji; https://imagej.nih.gov/ij/) ^92^ to convert image size from pixels to μm. Definition of regions of interest, subtraction of background, and fluorescence and area quantitation were performed Fiji ^92^.

### Statistical analysis

The data are presented as the mean ± standard error of the number of cells analyzed and measurements performed. Differences between two samples were assessed by t-test. Differences between four or more samples were assessed by one-way analysis of variance (ANOVA) followed by Tukey’s multiple comparison tests using GraphPad Prism version 7.01 software. *: 0.01<p<0.05; **: 0.001<p<0.01; ***: p<0.001.

### Supporting materials

Two supporting figures are available as supplementary data for this paper.

## Acknowledgements

The authors are grateful for the following institutions and people:

Synchrotron SOLEIL for beam time awarded at DISCO beamline (proposal 20160888) and all the support they benefited as SOLEIL’s users. Synchrotron ALBA for beam time awarded at MISTRAL beamline (proposals 2017022087, 2018022705) and all the support they benefited as ALBA’s users. FIOCRUZ Network of Technological Platforms for access to its facilities (Platforms RPT07C - Confocal and Electron Microscopy and RPT08L – Flow Cytometry). Dr. Bruna H. Marcon for support with the analyses performed at Platform RPT07C. Prof. Jeremy Mottram, Centre for Immunology and Infection, University of York, United Kingdom, for providing plasmids pGL2165 and pGL2166. Funding was provided by CAPES-COFECUB program (CAPES 862/2015, – COFECUB Me862-15), CNPq (NITZ: 312195/2015-0 and 304167/2019-3; BGG: 304027/2015-4 and 304788/2018-0), FIOCRUZ Inova program (3501948026).

## Author contributions

NITZ, BGG, FJ, and MR designed the experiments and the working plan. NITZ, FJ, GC, BGG, and MR collected and analyzed fluorescence data. FJ, EP, JJC and NITZ planned the experiments and prepared the samples for cryo-soft X-ray tomography. GC, FJ, EP, NITZ, BGG and PL collected and analyzed the cryo-soft X-ray tomography data. PMH and NITZ designed the PCR-base experiments for tagging of proteins. PMH and FRGC established the conditional strains and performed microscopic, flow cytometry and high content image analyses. GC and VR performed cell cultures. NITZ wrote the manuscript with input from other authors.

## Additional information

### Supplementary Information

The online version contains supplementary material.

### Data availability

Correspondence and requests for materials should be addressed to N.I.T.Z.

## References

1. Ohkura, H. et al. Cold-sensitive and caffeine-supersensitive mutants of the Schizosaccharomyces pombe dis genes implicated in sister chromatid separation during mitosis. EMBO J. 7, 1465–1473 (1988).

2. Kinoshita, N., Goebl, M. & Yanagida, M. The Fission Yeast dis3 + Gene Encodes a 110-kDa Essential Protein Implicated in Mitotic Control . Mol. Cell. Biol. 11, 5839–5847 (1991).

3. Mitchell, P., Petfalski, E., Shevchenko, A., Mann, M. & Tollervey, D. The exosome: A conserved eukaryotic RNA processing complex containing multiple 3’→5’ exoribonucleases. Cell 91, 457–466 (1997).

4. Clayton, C. et al. Genetic nomenclature for Trypanosoma and Leishmania. Mol. Biochem. Parasitol. 97, 221–224 (1998).

5. Dziembowski, A., Lorentzen, E., Conti, E. & Séraphin, B. A single subunit, Dis3, is essentially responsible for yeast exosome core activity. Nat. Struct. Mol. Biol. 14, 15–22 (2007).

6. Lebreton, A., Tomecki, R., Dziembowski, A. & Séraphin, B. Endonucleolytic RNA cleavage by a eukaryotic exosome. Nature 456, 993–996 (2008).

7. Schneider, C., Leung, E., Brown, J. & Tollervey, D. The N-terminal PIN domain of the exosome subunit Rrp44 harbors endonuclease activity and tethers Rrp44 to the yeast core exosome. Nucleic Acids Res. 37, 1127–1140 (2009).

8. Tomecki, R. et al. The human core exosome interacts with differentially localized processive RNases: HDIS3 and hDIS3L. EMBO J. 29, 2342–2357 (2010).

9. Schaeffer, D. et al. The exosome contains domains with specific endoribonuclease, exoribonuclease and cytoplasmic mRNA decay activities. Nat. Struct. Mol. Biol. 16, 56–62 (2009).

10. Kobyłecki, K., Drążkowska, K., Kuliński, T. M., Dziembowski, A. & Tomecki, R. Elimination of 01/A′-A0 pre-rRNA processing by-product in human cells involves cooperative action of two nuclear exosome-associated nucleases: RRP6 and Dis3. Rna 24, 1677–1692 (2018).

11. Allmang, C. et al. The yeast exosome and human PM-Scl are related complexes of 3’ → 5’ exonucleases. Genes Dev. 13, 2148–2158 (1999).

12. Makino, D. L., Baumgäerner, M. & Conti, E. Crystal structure of an RNA-bound 11-subunit eukaryotic exosome complex. Nature 495, 70–75 (2013).

13. Szczepińska, T. et al. DIS3 shapes the RNA polymerase II transcriptome in humans by degrading a variety of unwanted transcripts. Genome Res. 25, 1622–1633 (2015).

14. Anderson, J. S. J. & Parker, R. The 3’ to 5’ degradation of yeast mRNAs is a general mechanism for mRNA turnover that requires the SK12 DEVH box protein and 3’ to 5’ exonucleases of the exosome complex. EMBO J. 17, 1497–1506 (1998).

15. Pefanis, E. et al. RNA exosome-regulated long non-coding RNA transcription controls super-enhancer activity. Cell 161, 774–789 (2015).

16. Schneider, C., Kudla, G., Wlotzka, W., Tuck, A. & Tollervey, D. Transcriptome-wide Analysis of Exosome Targets. Mol. Cell 48, 422–433 (2012).

17. Estévez, A. M., Lehner, B., Sanderson, C. M., Ruppert, T. & Clayton, C. The roles of intersubunit interactions in exosome stability. J. Biol. Chem. 278, 34943–34951 (2003).

18. Cristodero, M., Böttcher, B., Diepholz, M., Scheffzek, K. & Clayton, C. The Leishmania tarentolae exosome: Purification and structural analysis by electron microscopy. Mol. Biochem. Parasitol. 159, 24–29 (2008).

19. Estévez, A. M., Kempf, T. & Clayton, C. The exosome of Trypanosoma brucei. EMBO J. 20, 3831–3839 (2001).

20. Guerra-Slompo, E. P., Cesaro, G., Guimarães, B. G. & Zanchin, N. I. T. Dissecting Trypanosoma brucei RRP44 function in the maturation of segmented ribosomal RNA using a regulated genetic complementation system. Nucleic Acids Res. 51, 396–419 (2023).

21. Cesaro, G. et al. Trypanosoma brucei RRP44: a versatile enzyme for processing structured and non-structured RNA substrates. Nucleic Acids Res. 51, 380–395 (2023).

22. Kumakura, N., Otsuki, H., Tsuzuki, M., Takeda, A. & Watanabe, Y. Arabidopsis AtRRP44A is the functional homolog of Rrp44/Dis3, an exosome component, is essential for viability and is required for RNA processing and degradation. PLoS One 8, e79219 (2013).

23. Zhang, W., Murphy, C. & Sieburth, L. E. Conserved RNaseII domain protein functions in cytoplasmic mRNA decay and suppresses Arabidopsis decapping mutant phenotypes. Proc. Natl. Acad. Sci. U. S. A. 107, 15981–15985 (2010).

24. Hou, D., Ruiz, M. & Andrulis, E. D. The ribonuclease Dis3 is an essential regulator of the developmental transcriptome. BMC Genomics 13, (2012).

25. Snee, M. J. et al. Collaborative control of cell cycle progression by the RNA exonuclease Dis3 and ras is conserved across species. Genetics 203, 749– 762 (2016).

26. Wu, D. & Dean, J. RNA exosome ribonuclease DIS3 degrades Pou6f1 to promote mouse pre-implantation cell differentiation. Cell Rep. 42, (2023).

27. Mukarami, H. et al. Ribonuclease activity of Dis3 is required for mitotic progression and provides a possible link between heterochromatin and kinetochore function. PLoS One 2, (2007).

28. Milbury, K. L. et al. Exonuclease domain mutants of yeast DIS3 display genome instability. Nucleus 10, 21–32 (2019).

29. Laffleur, B. et al. Noncoding RNA processing by DIS3 regulates chromosomal architecture and somatic hypermutation in B cells. Nat. Genet. 53, 230–242 (2021).

30. Laffleur, B. et al. RNA exosome drives early B cell development via noncoding RNA processing mechanisms. Sci. Immunol. 7, 1–14 (2022).

31. McIver, S. C. et al. The exosome complex establishes a barricade to erythroid maturation. Blood 124, 2285–2297 (2014).

32. Mehta, C., Fraga De Andrade, I., Matson, D. R., Dewey, C. N. & Bresnick, E. H. RNA-regulatory exosome complex confers cellular survival to promote erythropoiesis. Nucleic Acids Res. 49, 9007–9025 (2021).

33. Gudipati, R. K. et al. Extensive Degradation of RNA Precursors by the Exosome in Wild-Type Cells. Mol. Cell 48, 409–421 (2012).

34. Droll, D. et al. Disruption of the RNA exosome reveals the hidden face of the malaria parasite transcriptome. RNA Biol. 15, 1206–1214 (2018).

35. Laffleur, B. & Basu, U. Biology of RNA Surveillance in Development and Disease. Trends Cell Biol. 29, 428–445 (2019).

36. Wolin, S. L. & Maquat, L. E. Cellular RNA surveillance in health and disease. Science (80-.). 366, 822–827 (2019).

37. Gritti, I. et al. Loss of ribonuclease DIS3 hampers genome integrity in myeloma by disrupting DNA:RNA hybrid metabolism . EMBO J. 41, 1–22 (2022).

38. Davidson, L. et al. Rapid Depletion of DIS3, EXOSC10, or XRN2 Reveals the Immediate Impact of Exoribonucleolysis on Nuclear RNA Metabolism and Transcriptional Control. Cell Rep. 26, 2779–2791.e5 (2019).

39. Weißbach, S. et al. The molecular spectrum and clinical impact of DIS3 mutations in multiple myeloma. Br. J. Haematol. 169, 57–70 (2015).

40. Lionetti, M. et al. A compendium of DIS3 mutations and associated transcriptional signatures in plasma cell dyscrasias. Oncotarget 6, 26129–26141 (2015).

41. Segalla, S. et al. The ribonuclease DIS3 promotes let-7 miRNA maturation by degrading the pluripotency factor LIN28B mRNA. Nucleic Acids Res. 43, 5182– 5193 (2015).

42. Cesaro, G., Carneiro, F. R. G., Ávila, A. R., Zanchin, N. I. T. & Guimarães, B. G. Trypanosoma brucei RRP44 is involved in an early stage of large ribosomal subunit RNA maturation. RNA Biol. 16, 133–143 (2019).

43. Figarella, K. et al. Prostaglandin-induced programmed cell death in Trypanosoma brucei involves oxidative stress. Cell Death Differ. 13, 1802– 1814 (2006).

44. Uzcátegui, N. L. et al. Antiproliferative effect of dihydroxyacetone on Trypanosoma brucei bloodstream forms: Cell cycle progression, subcellular alterations, and cell death. Antimicrob. Agents Chemother. 51, 3960–3968 (2007).

45. Van Zandbergen, G., Lüder, C. G. K., Heussler, V. & Duszenko, M. Programmed cell death in unicellular parasites: A prerequisite for sustained infection? Trends Parasitol. 26, 477–483 (2010).

46. Barth, T. et al. Staurosporine-Induced Cell Death in Trypanosoma brucei and the Role of Endonuclease G during Apoptosis. Open J. Apoptosis 03, 16–31 (2014).

47. Boulon, S., Westman, B. J., Hutten, S., Boisvert, F. M. & Lamond, A. I. The Nucleolus under Stress. Mol. Cell 40, 216–227 (2010).

48. Pfister, A. S. Emerging role of the nucleolar stress response in autophagy. Front. Cell. Neurosci. 13, 1–18 (2019).

49. Li, F. J. et al. A role of autophagy in Trypanosoma brucei cell death. Cell. Microbiol. 14, 1242–1256 (2012).

50. Klionsky, D. J. et al. Guidelines for the use and interpretation of assays for monitoring autophagy. Autophagy 8, 445–544 (2012).

51. Proto, W. R., Jones, N. G., Coombs, G. H. & Mottram, J. C. Tracking autophagy during proliferation and differentiation of trypanosoma brucei. Microb. Cell 1, 9–20 (2014).

52. Hollville, E. & Martin, S. J. Measuring apoptosis by microscopy and flow cytometry. Curr. Protoc. Immunol. 2016, 14.38.1–14.38.24 (2016).

53. Siegel, T. N., Hekstra, D. R. & Cross, G. A. M. Analysis of the Trypanosoma brucei cell cycle by quantitative DAPI imaging. Mol. Biochem. Parasitol. 160, 171–174 (2008).

54. Levy, G. V. et al. Depletion of the SR-related protein TbRRM1 leads to cell cycle arrest and apoptosis-like death in Trypanosoma brucei. PLoS One 10, 1– 20 (2015).

55. Kobayashi, Y. et al. Nuclear swelling occurs during premature senescence mediated by MAP kinases in normal human fibroblasts. Biosci. Biotechnol. Biochem. 72, 1122–1125 (2008).

56. Filippi-Chiela, E. C. et al. Nuclear morphometric analysis (NMA): Screening of senescence, apoptosis and nuclear irregularities. PLoS One 7, (2012).

57. Yoon, K. B., Park, K. R., Kim, S. Y. & Han, S. Y. Induction of nuclear enlargement and senescence by sirtuin inhibitors in glioblastoma cells. Immune Netw. 16, 183–188 (2016).

58. Schneider, G. et al. Three-dimensional cellular ultrastructure resolved by X-ray microscopy. Nat. Methods 7, 985–987 (2010).

59. Groen, J., Conesa, J. J., Valcárcel, R. & Pereiro, E. The cellular landscape by cryo soft X-ray tomography. Biophys. Rev. 11, 611–619 (2019).

60. Moreno, S. N. J. & Docampo, R. The role of acidocalcisomes in parasitic protists. J. Eukaryot. Microbiol. 56, 208–213 (2009).

61. Docampo, R. The origin and evolution of the acidocalcisome and its interactions with other organelles. Mol. Biochem. Parasitol. 209, 3–9 (2016).

62. Sviben, S. et al. A vacuole-like compartment concentrates a disordered calcium phase in a key coccolithophorid alga. Nat. Commun. 7, 1–9 (2016).

63. Gal, A. et al. Native-state imaging of calcifying and noncalcifying microalgae reveals similarities in their calcium storage organelles. Proc. Natl. Acad. Sci. U. S. A. 115, 11000–11005 (2018).

64. McDermott, G., Le Gros, M. A., Knoechel, C. G., Uchida, M. & Larabell, C. A. Soft X-ray tomography and cryogenic light microscopy: the cool combination in cellular imaging. Trends Cell Biol. 19, 587–595 (2009).

65. Uchida, M. et al. Soft X-ray tomography of phenotypic switching and the cellular response to antifungal peptoids in Candida albicans. Proc. Natl. Acad. Sci. U. S. A. 106, 19375–19380 (2009).

66. Uchida, M. et al. Quantitative analysis of yeast internal architecture using soft X-ray tomography. Yeast 28, 227–236 (2011).

67. Hummel, E. et al. 3D Ultrastructural Organization of Whole Chlamydomonas reinhardtii Cells Studied by Nanoscale Soft X-Ray Tomography. PLoS One 7, (2012).

68. Welte, M. A. Nihms678421. 25, 1–24 (2016).

69. Olzmann, J. A. & Carvalho, P. Dynamics and functions of lipid droplets. Nat. Rev. Mol. Cell Biol. 20, 137–155 (2019).

70. Eckers, E. & Deponte, M. No Need for Labels: The Autofluorescence of Leishmania tarentolae Mitochondria and the Necessity of Negative Controls. PLoS One 7, 1–4 (2012).

71. Kunz, W. S. & Kunz, W. Contribution of different enzymes to flavoprotein fluorescence of isolated rat liver mitochondria. BBA - Gen. Subj. 841, 237–246 (1985).

72. Lienhart, W. D., Gudipati, V. & MacHeroux, P. The human flavoproteome. Arch. Biochem. Biophys. 535, 150–162 (2013).

73. Peikert, C. D. et al. Charting organellar importomes by quantitative mass spectrometry. Nat. Commun. 8, (2017).

74. Alexander, D. L., Schwartz, K. J., Balber, A. E. & Bangs, J. D. Developmentally regulated trafficking of the lysosomal membrane protein p67 in Trypanosoma brucei. J. Cell Sci. 115, 3253–3263 (2002).

75. Taylor, M. C., Mclatchie, A. P. & Kelly, J. M. Evidence that transport of iron from the lysosome to the cytosol in African trypanosomes is mediated by a mucolipin orthologue. Mol. Microbiol. 89, 420–432 (2013).

76. Schmidt, R. S. & Buẗikofer, P. Autophagy in Trypanosoma brucei: Amino acid requirement and regulation during different growth phases. PLoS One 9, 1–10 (2014).

77. Li, F. J. & He, C. Y. Acidocalcisome is required for autophagy in Trypanosoma brucei. Autophagy 10, 1978–1988 (2014).

78. Hope, R. et al. Transcriptome and proteome analyses and the role of atypical calpain protein and autophagy in the spliced leader silencing pathway in Trypanosoma brucei. Mol. Microbiol. 102, 1–21 (2016).

79. Goldshmidt, H. et al. Persistent ER stress induces the spliced leader RNA silencing pathway (SLS), leading to programmed cell death in Trypanosoma brucei. PLoS Pathog. 6, (2010).

80. Hope, R. et al. Phosphorylation of the TATA-binding protein activates the spliced leader silencing pathway in Trypanosoma brucei. Sci. Signal. 7, 1–9 (2014).

81. Proto, W. R., Coombs, G. H. & Mottram, J. C. Cell death in parasitic protozoa: Regulated or incidental? Nat. Rev. Microbiol. 11, 58–66 (2013).

82. Vandenabeele, P., Galluzzi, L., Vanden Berghe, T. & Kroemer, G. Molecular mechanisms of necroptosis: An ordered cellular explosion. Nat. Rev. Mol. Cell Biol. 11, 700–714 (2010).

83. Morlot, S. et al. Excessive rDNA Transcription Drives the Disruption in Nuclear Homeostasis during Entry into Senescence in Budding Yeast. Cell Rep. 28, 408–422.e4 (2019).

84. Fridy, P. C. et al. A robust pipeline for rapid production of versatile nanobody repertoires. Nat. Methods 11, 1253–1260 (2014).

85. Nunez, R. DNA measurement and cell cycle analysis by flow cytometry. Curr. Issues Mol. Biol. 3, 67–70 (2001).

86. Sorrentino, A. et al. MISTRAL: A transmission soft X-ray microscopy beamline for cryo nano-tomography of biological samples and magnetic domains imaging. J. Synchrotron Radiat. 22, 1112–1117 (2015).

87. Kremer, J. R., Mastronarde, D. N. & McIntosh, J. R. Computer Visualization of Three-dimensional Image Data Using IMOD. J. Struct. Biol. 116, 71–6 (1996).

88. Agulleiro, J. I. & Fernandez, J. J. Tomo3D 2.0 - exploitation of advanced vector eXtensions (AVX) for 3D reconstruction. J. Struct. Biol. 189, 147–152 (2015).

89. Schneider, C. A., Rasband, W. S. & Eliceiri, K. W. NIH Image to ImageJ: 25 years of image analysis. Nat. Methods 9, 671–675 (2012).

90. Belevich, I., Joensuu, M., Kumar, D., Vihinen, H. & Jokitalo, E. Microscopy Image Browser: A Platform for Segmentation and Analysis of Multidimensional Datasets. PLoS Biol. 14, 1–13 (2016).

91. Oberholzer, M., Morand, S., Kunz, S. & Seebeck, T. A vector series for rapid PCR-mediated C-terminal in situ tagging of Trypanosoma brucei genes. Mol. Biochem. Parasitol. 145, 117–120 (2006).

92. Schindelin, J., et al. Fiji: An open-source platform for biological-image analysis. Nat. Methods 9, 676–682 (2012).

