## Supplementary Figures S1 and S2 for "*Trypanosoma brucei* ultra-structure cell modifications caused by knockdown of the ribonuclease Rrp44/Dis3"

**This file includes:**

Figs. S1 to S2

### SUPPORTING FIGURES

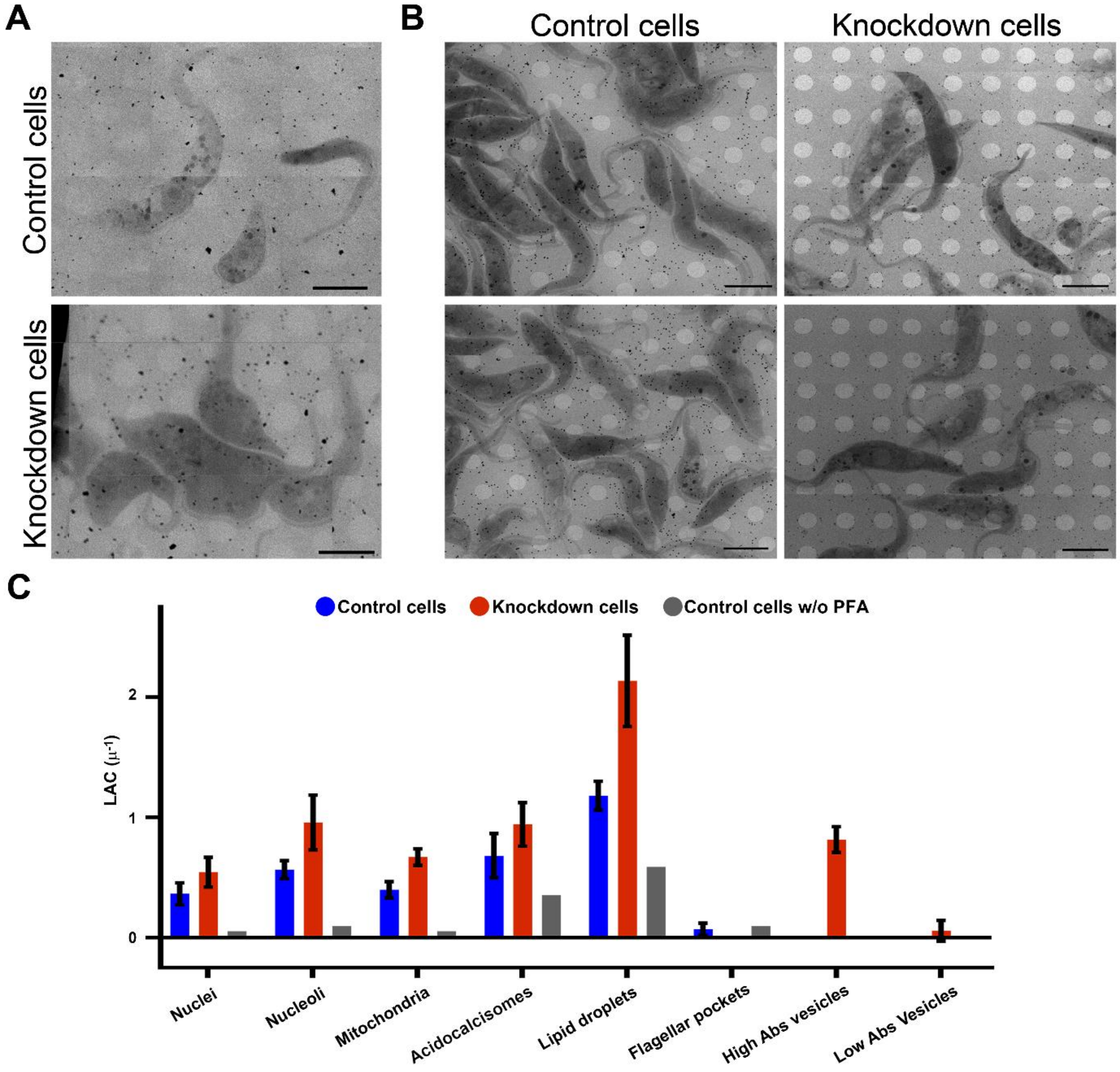


**Figure S1. Effect of PFA on the preservation of morphology and on linear absorption coefficient values.** (A) Representative X-ray mosaics of projections of cells not fixed with PFA. (B) Representative X-ray mosaics of projections used to select cells for cryo soft X-ray tomography. Scale bar, 5 µm. (C) Comparison of average linear absorption coefficient (LAC) for organelles in the *T. brucei* cells. Four to six tomograms (depending on the analyzed organelle) of PFA-fixed control and knockdown cells were used for quantification. Despite the difficulty to work with unfixed cell, we managed to reconstruct the three-dimensional (3D) structure of two control cells that have not been submitted to the PFA treatment. For comparison, average LAC for organelles of non-fixed control cells based on these two tomograms are also shown. Due to the low number, no error bars are shown for them. Although relatively short, the PFA treatment appears to have a high impact on LAC values. In PFA-treated control cells, the LAC of nuclei, nucleoli and mitochondria was approximately 5 times higher, while lipid droplets and acidocalcisomes showed LAC ~2 times higher when compared to the LAC of untreated control cells. In TbRrp44 knockdown cells, the LAC of these organelles was significantly higher than in control cells. However, it is not possible to know if this is a phenotype caused by TbRrp44 depletion, corresponding to a real increase in the LAC value or, if it is an artifact caused by PFA fixation.

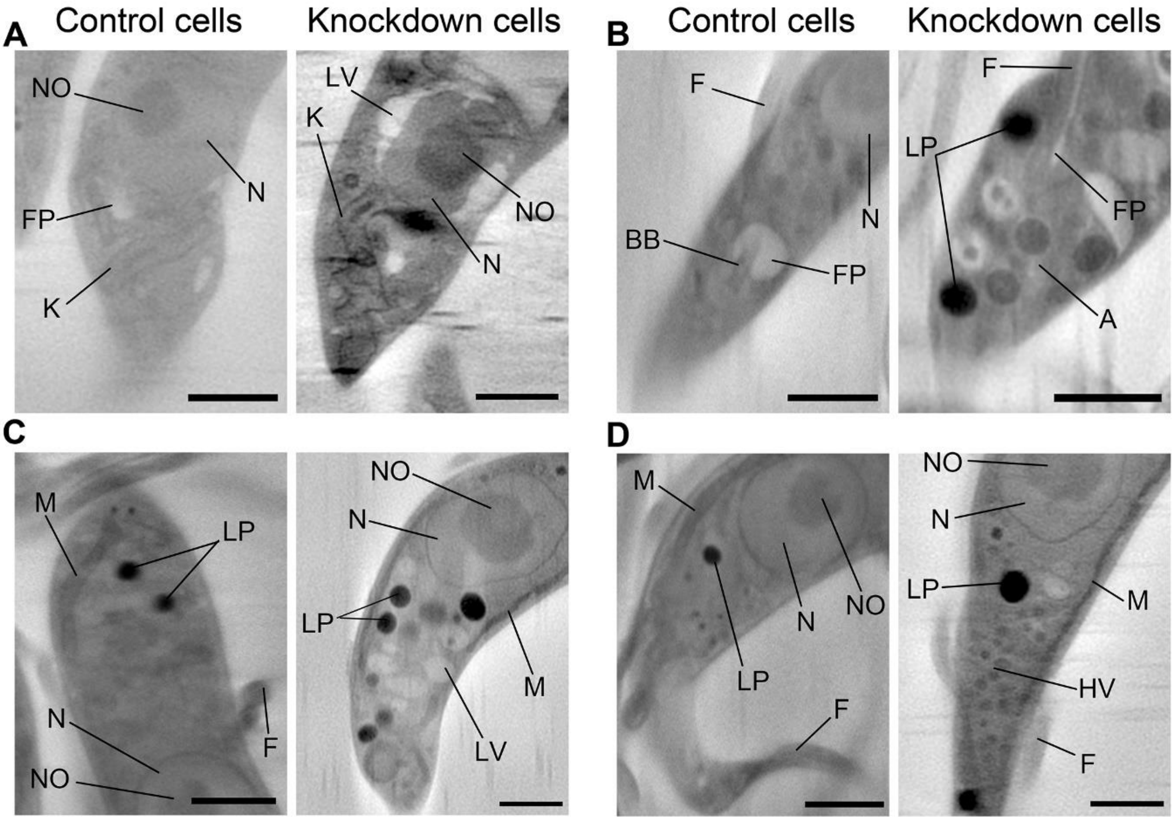


**Figure S2. Cryo-SXT comparison between control and TbRrp44 knockdown cells showing different alterations in the anterior and posterior cellular poles.** (A, B and C) Volume slices of the anterior pole showing that the kinetoplast remains intact in knockdown cells and a smaller and/or deformed flagellar pocket in comparison with the controls. (D) Volume slices of the posterior pole. Differences between control and knockdown cells include nuclei with different shapes, vacuoles around the nucleus, increase of the number of vesicles such as lipid droplets, high X-ray absorbing vesicles and low X-ray absorbing vacuoles. Letters indicate the organelles identified. N: nucleus; NO: nucleolus; K: kinetoplast; F: flagellum; M: mitochondrion; HV: high X-ray absorbing vesicles; LV: low X-ray absorption vacuoles; BB: basal body; A: acidocalcisomes; LP: lipid droplet. Scale bar, 1.5 µm.
